# Understanding How Actomyosin Dynamics Drive Apical Constriction by Systematically Tagging and Mapping Actin-Binding Proteins

**DOI:** 10.64898/2026.10.23.761371

**Authors:** Pu Zhang, Bailey N. de Jesus, Taylor N. Medwig-Kinney, David Q. Matus, Callista Yee, Theresa Gibney, Ariel Pani, Bob Goldstein

## Abstract

Apical constriction shapes tissues in fundamental events including gastrulation and neural tube formation. Diverse actin regulators play key roles in shaping cells, but the spatial distribution of most actin regulators in apical constriction is unknown. Here, we describe a systematic approach revealing the endogenous localization of 49 actin regulators in *C. elegans* gastrulation – nearly every actin-binding protein (ABP) expressed in the early embryo – and we use this atlas to test mechanistic hypotheses. We found that many actin regulators localized basolaterally, dependent on the Arp2/3 complex; that linear actin assembly factors enriched apically; and that apically constricting cells had an extra requirement for membrane-cortical linkers present in all cells. These findings answer some key questions about how ABPs function in apical constriction, and they establish a rich resource for investigating actin organization throughout early *C. elegans* development and a systematic *in vivo* protein localization reference for the actin cytoskeleton community.

## Introduction

Animals are composed in large part of epithelia that are bent into specific shapes. Apical constriction is one of the most common processes that bends epithelia, driving tissue invagination and cell extrusion during development and disease.^1^ Apical constriction is essential for gastrulation in many animals and neural tube formation in vertebrates, and failures in these processes can lead to severe congenital defects in humans such as spina bifida and anencephaly.^1,2^ The constriction of the apical side of a cell is powered by actomyosin contractility, with non-muscle myosin II (referred to as myosin here unless otherwise indicated) generating forces on networks of filamentous actin (F-actin) that are mechanically coupled to adherens junctions.^1,3,4^ While the force-generating role of myosin has been well characterized, how actin’s assembly, turnover, organization, and dynamic remodeling contribute to the mechanics and regulation of apical constriction remains poorly understood. Resolving this gap is critical for understanding how cytoskeletal architecture translates molecular-scale activity into cell-and tissue-scale force generation and morphogenesis.

Diverse F-actin networks are assembled and remodeled by a wide array of actin-binding proteins (ABPs) with specialized and often overlapping functions (summarized in Figure 1A).^5^ Actin nucleators such as the Arp2/3 complex and formins initiate branched and linear F-actin assembly, respectively.^6^ Capping protein binds barbed ends to regulate filament elongation and turnover, while debranching, depolymerizing, and severing proteins such as glia maturation factor (GMF), cofilin, and coronin promote actin turnover and network remodeling.^7^ F-actin can be further organized into higher-order structures by crosslinkers, bundlers, and scaffolding proteins including filamin and plastin.^8^ Myosins engage F-actin networks to generate contractile forces and mediate cargo transport.^9^ These actomyosin assemblies are coupled to membranes by linker proteins such as ezrin/radixin/moesin family (ERM) proteins, spectrins, and the cadherin-catenin complex, enabling force transmission and cell shape changes.^10–12^

**Figure 1.**
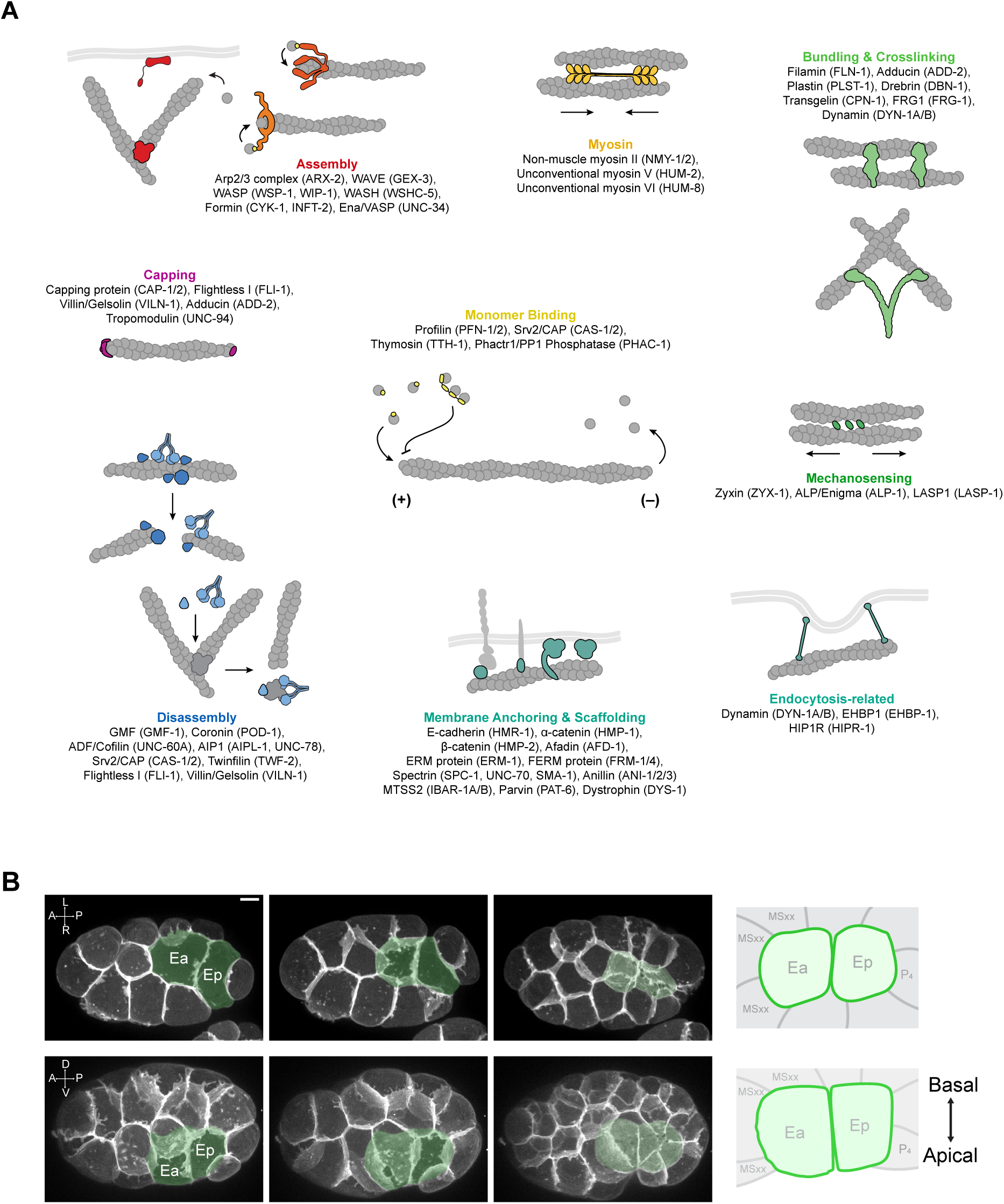
Overview of ABPs and *C. elegans* gastrulation. (A) Schematic representation of major functional classes of ABPs. Corresponding *C. elegans* proteins are indicated in parentheses. (B) Ventral (top) and lateral (bottom) views of *C. elegans* embryos expressing a membrane marker (mScarlet-I::PH) show internalization of the endodermal precursor cells Ea and Ep (pseudocolored) over time. Schematics to the right illustrate the relative positions of Ea, Ep, and neighboring cells and serve as references for ABP localization throughout the manuscript. The schematics were adapted from Zhang et al., 2025.^37^ Scale bar, 5 µm. See also Table S1.

The integrated roles of ABPs in the dynamic control of cell shape are perhaps best understood in migrating cells, where distinct combinations of ABPs assemble and remodel F-actin networks with specialized architectures and functions.^13^ At the leading edge, regulators including the Arp2/3 complex, capping protein, and cofilin work together to generate and remodel the dynamic branched actin network of lamellipodia that drives membrane protrusion.^14–16^ Along the cell body, stress fibers integrate actin assembly, filament crosslinking, and non-muscle myosin II activity to form contractile actomyosin bundles that contribute to cell adhesion and movement.^17^ Coordinated regulation among these machineries within a shared cytoplasm is essential for productive motion. For example, capping protein limits the elongation of individual filaments, while simultaneously promoting protrusion efficiency by competing with and liberating nucleation-promoting factors (NPFs) from barbed ends, stimulating Arp2/3 complex-dependent branched actin nucleation.^18^ Likewise, the front protrusion and rear retraction in migrating neutrophils are mediated by cell-scale actin retrograde flow, membrane tension, and dynamic myosin relocalization.^19^

The field’s understanding of apical constriction is only beginning to reach this level of mechanistic integration. Analogous to the situation for migrating cells, apical constriction may also rely on the regulated construction and remodeling of specialized F-actin networks by ABPs. Findings from our lab and others, using a variety of animal models, reveal that individual ABPs play important roles in apical constriction.^20–28^ These observations raise the possibility that multiple ABPs act in concert, coordinating F-actin network organization and movement during apical constriction.

To test this idea, and to advance our understanding of actin regulation in apical constriction toward the mechanistic depth achieved for cell migration, we decided to systematically examine the localization and dynamics of ABPs during gastrulation in the early *C. elegan*s embryo. Gastrulation in *C. elegans* begins at the 26-cell stage, when two endodermal precursor cells internalize from the surface to the embryo interior by apical constriction (Figure 1B). The combination of optical accessibility and powerful genetic tools makes *C. elegans* a tractable system for dissecting actin regulation on a large scale with cellular and subcellular resolution, in a complex, *in vivo* context.

We chose an imaging-first approach to understand actin regulation in apical constriction for three reasons. First, the subcellular localization of ABPs frequently correlates with and determines their molecular functions, enabling inferences about actin regulation to be drawn from their spatial distribution.^13,29^ Second, inherent to candidate-based loss-of-function approaches are a limitation of pleiotropy: several *C. elegans* ABPs are required for developmental events that precede endodermal precursor cell apical constriction; for example, depletion of formin CYK-1 or cofilin UNC-60A causes defects at the one-cell stage, necessitating analysis of their later roles by methods other than traditional loss of function experiments.^30,31^ Third, many ABPs are encoded by multiple homologs expressed in the early embryo, such that targeting individual proteins in classes of redundant contributors may yield false-negative results.

On this basis, we set out to generate a comprehensive, endogenous fluorescent protein tagging resource to visualize essentially the complete set of ABPs expressed in early *C. elegans* embryos. Using a combination of unbiased and targeted strategies, we identified and tagged (or used existing strains where available) 49 ABPs. Tags were inserted into endogenous loci so that 100% of each protein population would be tagged. We rigorously validated whether tagging disrupted protein functions, re-tagging at different sites where necessary. Visualizing the localization of all of the classes of ABPs provided a panoramic view of actin regulatory proteins in the gastrulating embryo and suggested potential functional hubs, which we further investigated through colocalization analyses and RNAi perturbations. This work established an integrated framework for linking actin regulators to local functions in a developing embryo, uncovered new principles of spatiotemporal actin control during morphogenesis, and created a broadly applicable resource for dissecting cytoskeletal organization in a model organism *in vivo*.

## Results

### Identification and endogenous tagging of ABPs expressed during endodermal precursor cell apical constriction

To comprehensively identify all ABPs expressed during apical constriction, we first collected *C. elegans* genes annotated with the Gene Ontology (GO) term “actin-binding” (GO:0003779) and organized them according to known molecular functions. We supplemented this list by searching the literature for ABP classes described in other systems that were absent or incompletely represented in the *C. elegans* GO annotation. We then examined mRNA expression from these genes using our publicly available single-cell RNA-sequencing dataset spanning the 1-cell stage through formation of the endodermal precursor cells.^32^ Based on previous empirical experience, we selected genes with a median mRNA abundance ≥ 20 RPKM (reads per kilobase of transcript per million mapped reads) in endodermal precursor cells or in any of their precursor lineages (P_0_, P_1_, EMS, and E). We also included additional genes with lower expression when previous studies or known molecular functions suggested potential relevance. For some protein complexes, representative subunits were selected for analysis. The resulting set of candidate ABPs was organized by functional class (Table S1; Figure 1A).

To determine the subcellular localization of ABPs expressed in the early embryo, we next assembled a collection of endogenously tagged strains for these proteins. We generated new endogenously tagged alleles for 37 genes and obtained strains generated by other laboratories for the remaining genes. All newly generated and acquired strains included in the analysis were sequence-verified at the edited locus. Because endogenous tagging can alter protein function or localization, we systematically evaluated the functionality of each tagged allele before interpreting its localization. For essential genes, we considered 99-100% embryonic viability as evidence that tagging did not perturb protein function; exceptions are explicitly noted and considered when interpreting the localization data. For other genes, we assessed hatching rates, plate-level phenotypes, and localization in embryos and adults in relation to previously reported functional and localization data from *C. elegans* and other organisms. For all genes, we noted any discrepancies between our observations and previous reports (Table 1). This validation showed that the large majority of ABPs could be fluorescently tagged without detectable disruption of function. Strain design, validation results, hatching rates, and comparisons with previous studies are summarized in Tables S2 and S3, with gene-specific information accompanying the localization data in Figure 2 and Table 1.

**Figure 2.**
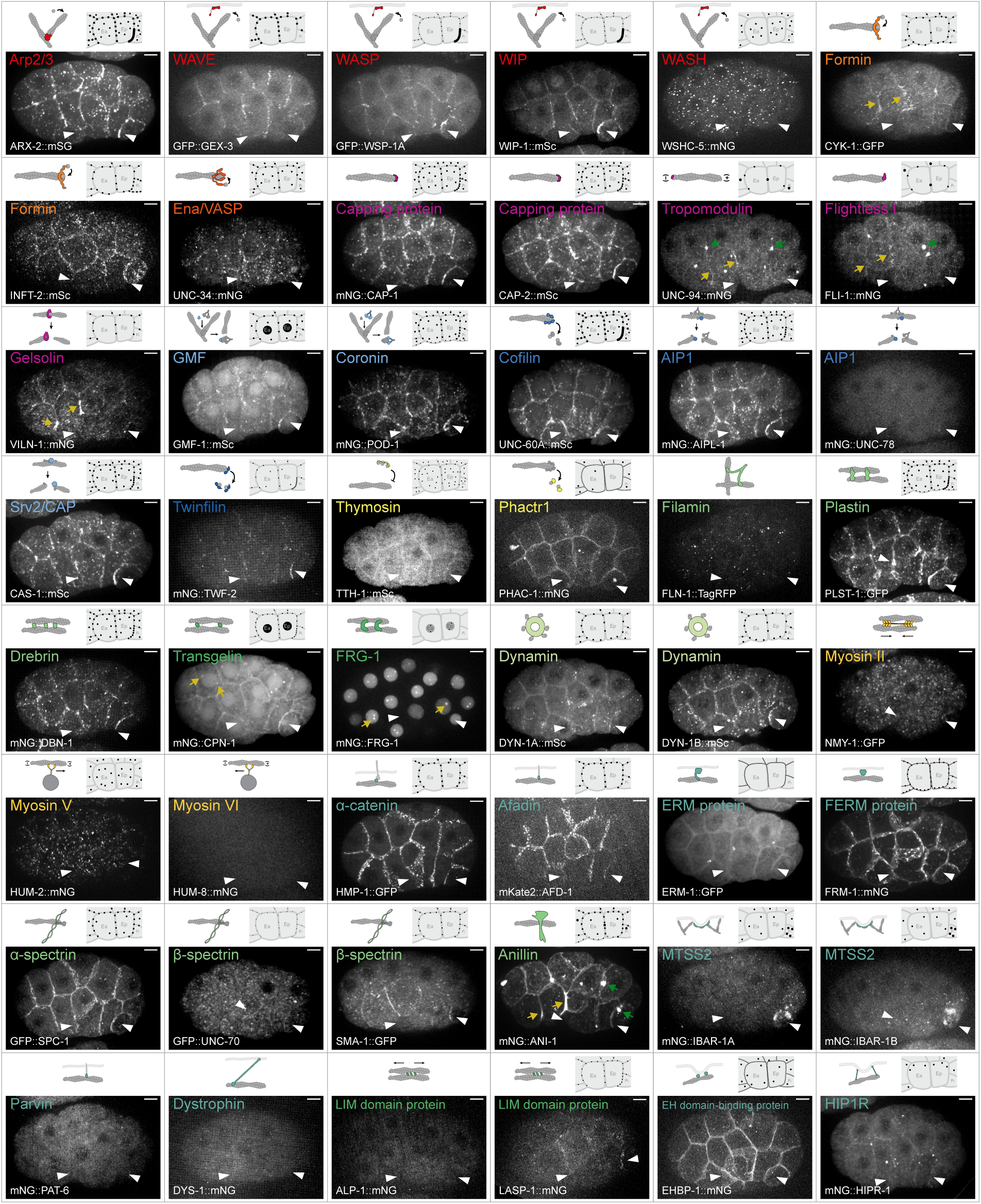
Localization patterns of ABPs during gastrulation. Localization patterns of 48 ABPs. For each ABP, the upper left shows a diagram depicting its interactions with G-actin and/or F-actin; the upper right shows a schematic summarizing its major localization patterns in Ea, Ep, and neighboring cells; and the bottom shows a maximum-intensity projection of an embryo expressing the corresponding fluorescently tagged ABP. White arrowheads indicate Ea and Ep. Yellow and green arrows highlight localization features described in Table 1. Scale bars, 5 µm. See also Figures S1-S2 and Tables S2-S4.

**Table 1.**
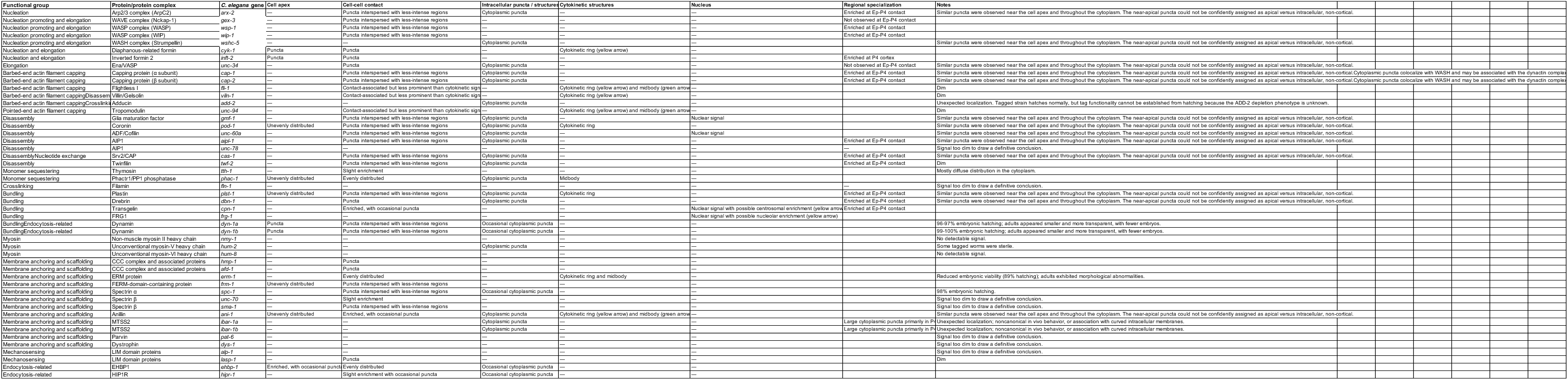
Localization patterns of ABPs during *C. elegans* gastrulation. Localization of endogenously tagged ABPs was scored at the apical cortex, cell-cell contacts, intracellular puncta or structures, cytokinetic structures, and nucleus. Region-specific patterns are listed under Regional specialization. An em dash (—) indicates that the indicated localization was not observed under the imaging conditions used. Protein-specific observations and limitations are provided in the Notes column.

Together, these efforts generated a sequence-verified and functionally evaluated collection of endogenously tagged ABPs to enable systematic comparison of their spatial organization.

### A localization atlas reveals spatial convergence and specialization of actin-binding proteins

We next filmed embryos of each strain by spinning disk confocal microscopy. During this analysis, we found some N-and C-terminally mNeonGreen-tagged proteins showed unexpected cytoplasmic patterns resembling mitochondrial localization (Figure S1, Table S4). Among these, we noticed an obvious phenotypic defect in UNC-60A::mNG animals: homozygous tagged animals produced no viable progeny. To test if mNeonGreen can recruit a small subset of tagged proteins to incorrect sites, we generated alternative endogenously tagged alleles by changing the fluorescent protein and/or tag position (Figure S1, Table S4). Altering the tag configuration shifted proteins from this prominent mitochondria-like localization to distributions more consistent with previously reported localization or known protein function (Figure S1, Table S4), and these strains were used for localization analyses and mechanistic experiments.

We then systematically examined the localization of the endogenous ABPs in gastrulating embryos. Part of our goal was to define the key subcellular landmarks where F-actin may be regulated in distinct ways, akin to the leading edge, focal adhesions, stress fibers, and trailing edge of crawling cells. For each protein, we scored localization at defined cellular regions and structures, including the cell apex, cell-cell contacts, intracellular puncta or structures, cytokinetic structures, and the nucleus. We also documented narrower patterns, such as protein enrichment at the contact between the endodermal precursor Ep and the germline precursor P_4_, when observed. The resulting localization profiles are summarized in Table 1. Figure 2 presents the corresponding visual atlas, with representative fluorescence images for each protein together with diagrams illustrating the observed localization patterns. We interpret failure to detect a protein at a particular cellular region as indicating that the protein may either be absent or present at levels below the detection limit of our imaging. Potential limitations associated with individual tagged alleles are noted in Table 1.

Considering the atlas as a whole revealed prominent features of ABP organization that are expanded upon below: functionally diverse actin regulators converged at common cellular regions but often adopted distinct spatial organizations within those regions, and proteins with closely related biochemical activities frequently differed in their spatial distributions (Table 1; Figure 2).

### Diverse actin regulators converge at common cellular and subcellular regions while retaining distinct spatial organizations

We found that the most widespread pattern observed was the localization of functionally diverse ABPs at sites of cell-cell contact between all or most cells, suggesting significant regulation of actin filaments in the basolateral cell cortex (Table 1; Figure 2). These cell contact-associated proteins are predicted to regulate actin in diverse ways; they include proteins implicated in other systems in actin assembly, filament-end regulation and turnover, filament bundling and crosslinking, and membrane-cortex association. For example, the Arp2/3 complex protein ARX-2, formins CYK-1 and INFT-2, and Ena/VASP UNC-34 represent distinct modes of actin assembly or elongation (see Figure 1 for protein homology classes) yet were all detected at cell-cell contacts. Proteins associated with filament turnover, including glia maturation factor GMF-1, cofilin UNC-60A, AIP1 AIPL-1, Srv2/CAP CAS-1, and twinfilin TWF-2, also accumulated near cell contacts, as did higher-order actin organizers such as plastin PLST-1, drebrin DBN-1, transgelin CPN-1, and anillin ANI-1 and membrane-associated proteins including ERM protein ERM-1, FERM-domain-containing protein FRM-1, and spectrin SPC-1. Thus, cell-cell contacts emerge from the atlas as major localization sites at which diverse actin-regulatory proteins converge during gastrulation.

Despite their shared localization at cell-cell contacts, these proteins exhibited distinct spatial distributions within this region. Here, we focused specifically on their distributions at cell-cell contacts, although many of these proteins also localized to other cellular regions (Table 1; Figure 2). A substantial set of proteins, including Arp2/3 complex ARX-2, capping protein CAP-1, glia maturation factor GMF-1, coronin POD-1, cofilin UNC-60A, AIP1 AIPL-1, Srv2/CAP CAS-1, twinfilin TWF-2, plastin PLST-1, FERM-domain-containing protein FRM-1, and spectrin SPC-1, showed prominent puncta along cell-cell contacts together with less intense, nonpunctate signal along the contacts. Other proteins, including formin CYK-1, formin INFT-2, Ena/VASP UNC-34, drebrin DBN-1, and alpha-catenin HMP-1, localized predominantly to discrete puncta at cell-cell contacts, with little detectable signal between puncta. Proteins such as transgelin CPN-1 and anillin ANI-1 showed broader contact enrichment with few puncta. Thus, cell-cell contacts contained multiple spatial organizations of actin regulators. The recurrence of similar patterns among proteins with different predicted biochemical activities further raised the possibility that selected ABPs may be organized into shared, contact-associated puncta, a possibility examined mechanistically below.

Distinct spatial organizations were also apparent at the cell apex and cytokinetic structures. At the cell apex, formins CYK-1 and INFT-2 and dynamin DYN-1A/B had punctate distributions. In contrast, coronin POD-1, plastin PLST-1, FERM-domain-containing protein FRM-1, anillin ANI-1, and Phactr1/PP1 phosphatase PHAC-1 were distributed unevenly, whereas EH domain-binding protein EHBP-1 was distributed more uniformly (Figure 2, Figure S2B). Apical localization is not always apparent in maximum-intensity projections of 8-10 optical sections; Figure S2B presents selected optical sections showing apical localization either at the apical surface (in the apical-most section or a maximum-intensity projection of the two apical-most sections) or along cell edges (in a middle section). Several proteins also localized to cytokinetic structures – to the cytokinetic ring, midbody, or both (Table 1; Figure 2, yellow and green arrows). Together, these observations show that some ABPs converge to the same general cellular region while retaining some distinct spatial organizations.

The atlas also revealed non-cortical ABP populations in the cytoplasm or nucleus. Multiple proteins formed cytoplasmic puncta, and several displayed nuclear signal during interphase. Glia maturation factor GMF-1 and cofilin UNC-60A showed nuclear signal in addition to their contact-associated distributions, whereas transgelin CPN-1 showed nuclear signal and possible centrosomal enrichment, and FRG-1 showed nuclear signal and enrichment in spots reminiscent of nucleoli.^33^ Other proteins exhibited unexpected cytoplasmic patterns, including large IBAR-1A/B-positive cytoplasmic puncta observed primarily in the germline founder cell, P_4_. Although localization alone does not establish the functions of these intracellular pools, these observations identify additional spatially restricted ABP populations for future investigation.

### Proteins with related biochemical activities diverge spatially, and cell-cell interfaces differ in ABP enrichment

Comparison of functionally similar proteins revealed substantial spatial divergence. The two formins CYK-1 and INFT-2, for example, both formed puncta at the apical cortex and cell-cell contacts, but their broader distributions differed (Figure 2, Figure S2B). CYK-1 was prominently associated with the cytokinetic ring as has been seen before,^30,34^ whereas INFT-2 showed no comparable cytokinetic enrichment and instead was enriched at the P_4_ cell cortex (Figure 2, Figure S2B). Thus, even proteins predicted to promote similar classes of actin assembly can be deployed to different actin-rich structures, consistent with the differential utilization and sorting of ABPs among distinct F-actin networks in diverse systems.^35,36^

Differences were also apparent among capping proteins. Capping protein CAP-1 showed a prominent punctate distribution at cell-cell contacts, whereas flightless I FLI-1 and villin VILN-1 were more prominently associated with cytokinetic structures and had weaker contact-associated signals. The pointed-end capping protein UNC-94 similarly showed prominent cytokinetic ring and midbody localization. These patterns identify distinct spatial biases among filament-end regulators and suggest that proteins with related effects on filament dynamics may participate in different F-actin networks.

Actin bundling proteins likewise showed both shared and distinct localization patterns. Plastin PLST-1, drebrin DBN-1, and transgelin CPN-1 were all associated with cell-cell contacts and were enriched at the Ep-P_4_ interface (the interface between an endoderm precursor and the germline founder cell), but their broader localization patterns differed. Plastin PLST-1 additionally displayed an uneven apical pattern and localized to the cytokinetic ring, drebrin DBN-1 formed discrete contact and cytoplasmic puncta, and transgelin CPN-1 showed broader contact enrichment with occasional puncta together with nuclear signal. These examples illustrate that shared biochemical activities or localization to a common cellular region do not necessarily correspond to identical spatial deployment.

Another example of regional specialization emerged from comparison of individual cell-cell interfaces. A distinct group of proteins, including Arp2/3 complex ARX-2, capping protein CAP-1, AIP1 AIPL-1, Srv2/CAP CAS-1, twinfilin TWF-2, plastin PLST-1, drebrin DBN-1, and transgelin CPN-1, was enriched at the Ep-P_4_ contact. In contrast, Ena/VASP UNC-34, despite its punctate localization to other cell-cell contacts, was not observed at the Ep-P_4_ contact. Formin INFT-2 showed a different form of regional asymmetry, with enrichment in puncta broadly throughout the P_4_ cell cortex rather than specifically at the Ep-P_4_ contact. These differences demonstrate that neighboring cortical and cell-cell contact regions can contain distinct actin regulators, and they highlight additional landmarks populated with distinct sets of actin regulators.

Together, the atlas reveals an organization in which diverse actin-regulatory activities converge within common cellular territories while individual proteins retain distinct subcellular and regional distributions. This combination of spatial convergence and specialization provides a framework for generating specific hypotheses about how the actin cortex is organized to drive morphogenesis. We therefore asked whether recurrent localization patterns reflect shared mechanisms of ABP organization, whether differences between cell-cell interfaces reveal new regulatory relationships, and whether distinct spatial organizations of ABPs could uncover complementary mechanisms that enable the cortex to withstand and transmit forces during apical constriction.

### Arp2/3 contributes to the organization of diverse actin regulators at cell-cell contacts

Having established a spatiotemporal map of ABP localization during endodermal precursor cell internalization, we next sought to use it to address some specific hypotheses. Because the localization of many different ABPs near sites of cell-cell contact was unexpected, we first asked whether this recurring localization pattern might reflect shared mechanisms of spatial organization. Most prominent among the patterns we observed were localization patterns resembling that of the Arp2/3 complex subunit ARX-2, including punctate-interspersed localization basolaterally at cell-cell contacts and cytoplasmic puncta. These proteins included the barbed-end capping protein CAP-1, multiple actin-turnover regulators, and actin-bundling proteins (Figure 2, 3A and S4A). Quantitative analysis of representative candidates confirmed co-localization with ARX-2 across the subcellular regions examined (Figure 3C and S4C).

**Figure 3.**
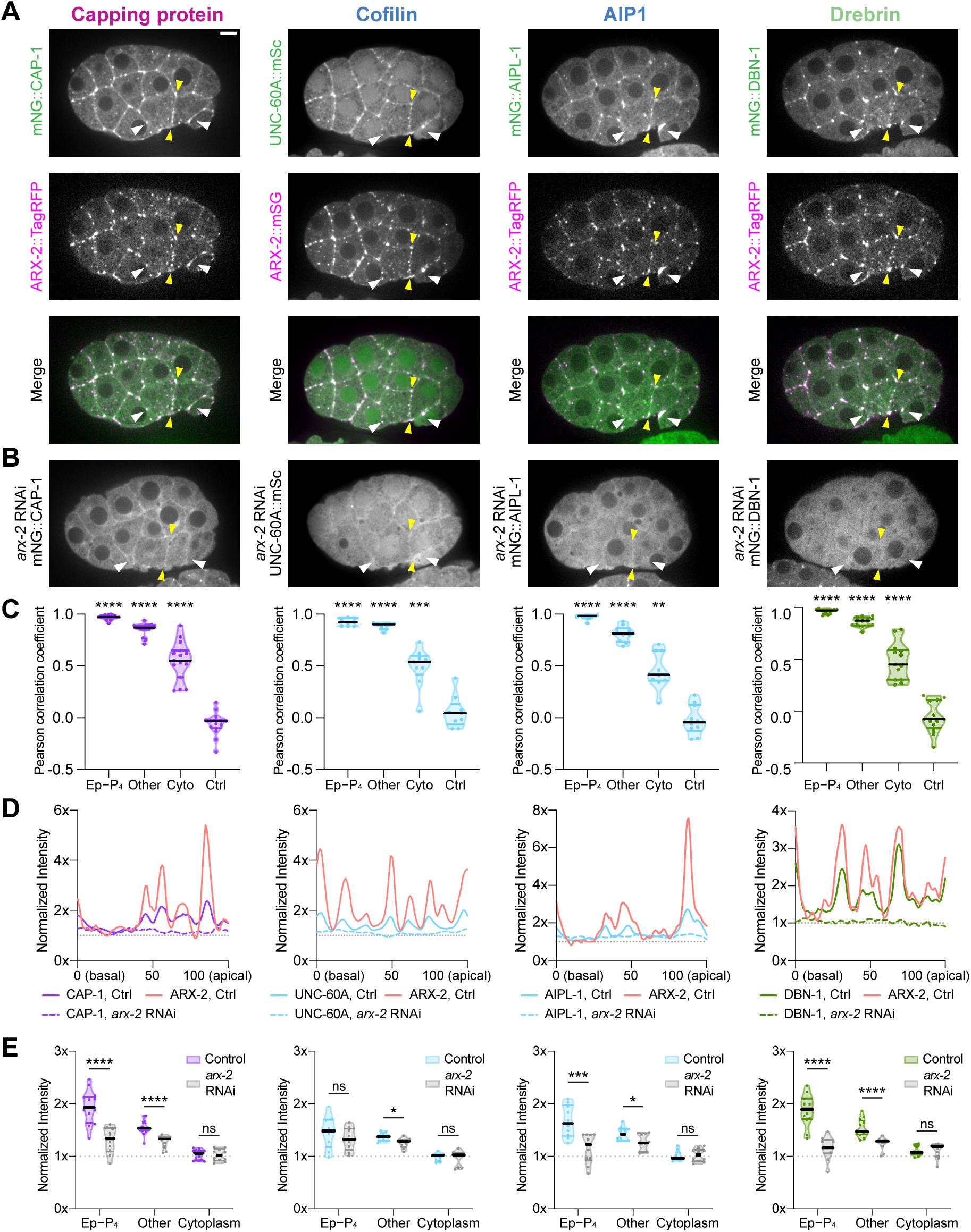
Arp2/3 contributes to the organization of diverse actin regulators at cell-cell contacts. (A) Co-visualization of CAP-1::mNG, UNC-60A::mSc, mNG::AIPL-1, and mNG::DBN-1 with ARX-2::TagRFP/mSG. (B) Localization of CAP-1::mNG, UNC-60A::mSc, mNG::AIPL-1, and mNG::DBN-1 following *arx-2* RNAi. (C) Quantification of colocalization between each protein of interest and ARX-2. Violin plots show Pearson correlation coefficients measured at the Ep-P_4_ contact, other cell-cell contacts, the cytoplasm, and a negative control generated by rotating one channel 180°. PCCs were Fisher *r*-to-*z* transformed before statistical analysis. (D) Representative intensity profiles along the Ea-Ep contact in control and *arx-2* RNAi-treated embryos. Colors indicate proteins; solid and dashed lines indicate control and *arx-2* RNAi conditions, respectively. (E) Quantification of protein localization at the Ep-P_4_ contact, other cell-cell contacts, and the cytoplasm in control and *arx-2* RNAi-treated embryos. For (C), Fisher-transformed PCCs were analyzed using repeated-measures one-way ANOVA with Geisser-Greenhouse correction followed by Dunnett’s multiple-comparisons test comparing each cellular region with the negative control. For (E), comparisons between control and *arx-2* RNAi were performed using an unpaired t test, an unpaired t test with Welch’s correction, or a Mann-Whitney test based on data distribution and variance. White arrowheads indicate Ea and Ep; yellow arrowheads indicate the Ea-Ep contact. *P < 0.05, **P < 0.01, ***P < 0.001, ****P < 0.0001. Scale bars, 5 μm. See also Figure S2 and S3.

ARX-2 and these colocalizing ABPs were particularly enriched in discrete puncta along cell-cell contacts. Because our previous work showed that contact-associated, rather than vesicle-associated, Arp2/3 contributes to apical constriction, we focused on this subcellular population.^37^ Along the Ea-Ep interface (Figure 3A and S4A, yellow arrowheads), ARX-2 and each candidate displayed coincident intensity peaks separated by regions of lower signal, indicating enrichment at similar positions along the contact (Figure 3D and S4E, solid lines). As a control, we examined the cadherin HMR-1, which localizes to cell-cell contacts but lacks the cytoplasmic punctate pattern observed for ARX-2 and the selected ABPs.^38^ HMR-1 showed lower colocalization with ARX-2 than did the selected ABPs (Figure S4D), as well as less concordant intensity profiles along the Ea-Ep interface (Figure S4E). Together, these results indicate that the coincident enrichment of ARX-2 and the selected ABPs reflects spatial organization within cell-cell contacts rather than their shared localization to the contact region.

The proteins that colocalized with ARX-2 are predicted by homology to regulate distinct steps of F-actin network assembly and remodeling, including barbed-end capping, filament turnover, and bundling. Their coincident enrichment therefore raised the possibility that these puncta represent hubs of co-localized proteins that function together, much as the proteins of a focal adhesion do in crawling cells, for example.^39^ Such organization could arise through independent recruitment by a common upstream cue or through Arp2/3-generated F-actin networks that recruit or retain other ABPs.

To test whether Arp2/3 contributes to the spatial organization of these contact-associated ABPs, we depleted ARX-2 by injection of *arx-2* dsRNA synthesized from *C. elegans* cDNA. We assessed knockdown efficiency by imaging unlabeled wild-type embryos, ARX-2-tagged embryos, and ARX-2-tagged embryos following *arx-2* RNAi side by side under identical conditions. RNAi reduced ARX-2 fluorescence to approximately 2% of the untreated tagged control, approaching the unlabeled wild-type background level (Figure S3A).The same validation strategy was used for subsequent RNAi experiments targeting other proteins in this study.

ARX-2 depletion reduced contact-associated enrichment of all candidates tested (Figures 3E and S4F) and eliminated their discrete intensity peaks, producing more uniform profiles along the Ea-Ep interface (Figure 3D and S4E, dashed lines). The magnitude of these effects varied, and some proteins, such as cofilin UNC-60A and coronin POD-1, retained residual contact-associated localization, indicating that their localization is not exclusively Arp2/3-dependent (Figures 3B, 3E, S4B, and S4F). Nevertheless, the shared loss of punctate organization supports a role for Arp2/3 in establishing this spatial pattern.

Together, these results indicate that Arp2/3 contributes to the spatial organization of functionally distinct ABPs into discrete domains at cell-cell contacts. These experiments do not distinguish direct molecular recruitment from indirect dependence on the Arp2/3-generated F-actin network, but they demonstrate that the recurrent contact pattern identified by the atlas reflects a shared Arp2/3-dependent organization. By concentrating regulators of capping, turnover, and bundling at common cortical sites, these domains may provide a platform for coordinating multiple aspects of F-actin-network remodeling during apical constriction. The results imply that Arp2/3-enriched puncta in the basolateral cell cortex, which we have implicated functionally in apical constriction before,^37^ are landmarks where multiple F-actin regulators localize, much like the leading edge of crawling cells.^13^

### The apical cortex is enriched with only linear nucleators during apical constriction, and the basolateral cortex has both branched and linear nucleators

The atlas revealed spatial differences among proteins that promote actin assembly. While our previous work showed that Arp2/3 is enriched primarily at basolateral cell-cell contacts rather than the apical cortex,^37^ here we found that the formins CYK-1 and INFT-2 localized to cell-cell contacts and, in contrast to Arp2/3, also formed puncta at the apical cortex (Figure 2, Figure S2B). This result identifies formins as the only known actin assembly factors prominently present in the constricting apical cortex.

This result left us uncertain about where Ena/Vasp UNC-34 functions, for example either apically together with the formins, or basolaterally with Arp2/3. Ena/Vasp UNC-34 can promote actin assembly by elongating growing filament barbed ends^40^ and could therefore act on filaments generated by either branched-or linear-actin nucleators. Although the relatively low abundance of UNC-34 limited our ability to define its cortical distribution, we could detect it at cell-cell contacts, where it formed puncta, but it was not observed at the Ep-P_4_ contact (Figure 2 and Figure 4A). This distribution was notable given our previous finding that distinct NPFs regulate Arp2/3 regionally during gastrulation. WAVE promotes Arp2/3 localization at most cell-cell contacts but is largely absent from the Ep-P_4_ interface, whereas WASP is preferentially associated with Ep-P_4_ and regulates Arp2/3 at this interface.^37^ The similarity between the spatial distributions of UNC-34 and WAVE therefore suggested that UNC-34 preferentially function with WAVE-regulated Arp2/3 networks.

**Figure 4.**
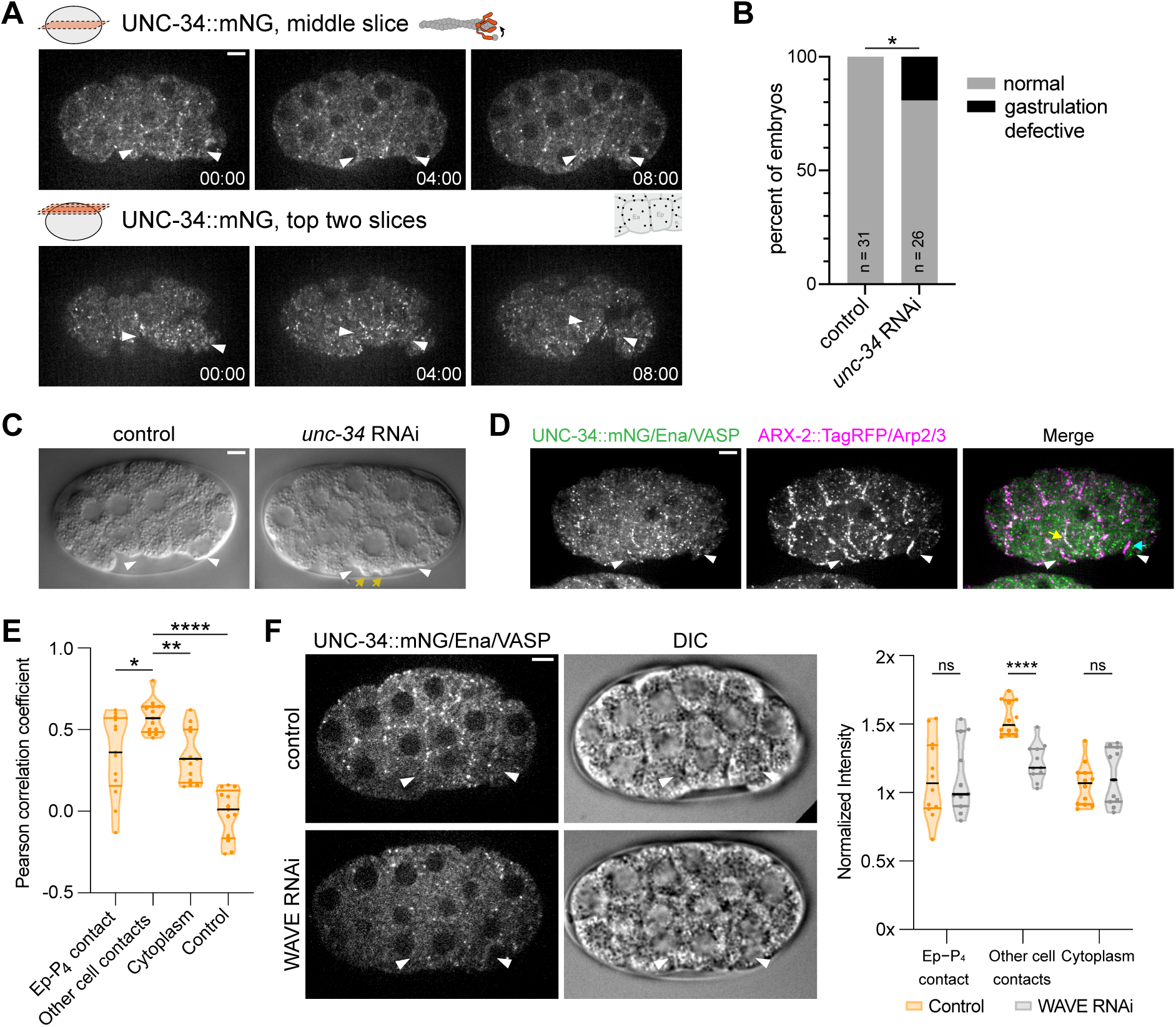
UNC-34/Ena/VASP cooperates with WAVE-Arp2/3 at a subset of cell-cell contacts. (A) Localization of UNC-34::mNG during gastrulation. Top, optical section through the middle of the embryo; bottom, maximum-intensity projection of the two apical-most optical sections. (B) Quantification of the effects of *unc-34* RNAi on gastrulation. (C) Micrographs from time-lapse DIC movies of control and *unc-34* RNAi embryos. Yellow arrows indicate blebs in the *unc-34* RNAi embryo. (D) Covisualization of UNC-34::mNG and ARX-2::TagRFP (E) Quantification of colocalization between UNC-34 and ARX-2. Violin plots show Pearson correlation coefficients measured at Ep-P_4_ contacts, other cell-cell contacts, the cytoplasm, and a negative-control where one channel was rotated 180°. PCCs were Fisher *r*-to-*z* transformed before statistical analysis. (F) Left, UNC-34::mNG localization in control and WAVE RNAi-treated embryos. Right, quantification of UNC-34 enrichment at the Ep-P_4_ contact, other cell-cell contacts, and the cytoplasm in control and WAVE RNAi-treated embryos. For categorical data in (B), statistical significance was assessed using Fisher’s exact test. For (E), Fisher-transformed PCCs were analyzed using repeated-measures one-way ANOVA with Geisser-Greenhouse correction followed by Dunnett’s multiple-comparisons test. For (F), comparisons were performed using unpaired t tests. White arrowheads indicate Ea and Ep. *P < 0.05, **P < 0.01, ****P < 0.0001. Scale bars, 5 μm. See also Figure S2.

Ena/VASP is known already to have context-dependent roles in apical constriction – it disrupts apical constriction when ectopically enriched in *Drosophila* ventral furrow cells but promotes apical F-actin accumulation and constriction during *Xenopus* neurulation and mouse lens placode invagination.^41–44^ Thus, Ena/VASP can either antagonize or promote apical constriction depending on developmental context. In *C. elegans*, UNC-34 can cooperate with WAVE/WASP-Arp2/3 during morphogenesis and migration, but can also oppose WASP-Arp2/3 during axon guidance.^45–48^ These findings suggest that UNC-34 can cooperate with or oppose WAVE/WASP-Arp2/3 pathways depending on cellular context.

To test whether UNC-34 contributes to *C. elegans* apical constriction, we depleted UNC-34 by RNAi and confirmed that UNC-34::mNG fluorescence was reduced to approximately 1% of the untreated tagged control (Figure S3B). Depletion caused mild gastrulation defects in some embryos (Figure 4B) and, notably, induced membrane blebs that were not observed in wild-type embryos (Figure 4C, yellow arrows). Similar blebbing occurs after Arp2/3 or WAVE depletion.^21,49^ These phenotypic similarities suggest that UNC-34 may cooperate with WAVE-Arp2/3 to maintain cortical integrity during constriction.

To test this idea, we generated embryos co-expressing UNC-34::mNG and ARX-2::TagRFP. The proteins colocalized at cell-cell contacts (Figure 4D, yellow arrow), but showed little overlap at the Ep/P_4_ boundary (Figure 4D, cyan arrow), where WASP, but not WAVE, is strongly enriched.^37^ Quantitative colocalization analysis supported this regional difference: UNC-34/Ena/VASP showed greater colocalization with Arp2/3 at non-Ep/P_4_ cell-cell contacts than in other regions (Figure 4E).

To test whether WAVE contributes to this organization, we examined UNC-34 localization following depletion of GEX-3, a WAVE complex subunit (hereafter referred to as WAVE RNAi). We validated previously the efficiency of WAVE RNAi.^37^ WAVE RNAi reduced the enrichment of UNC-34::mNG at non-Ep/P_4_ contacts (Figure 4F), indicating that WAVE is required for UNC-34 accumulation at these sites.

Together, these results identify a previously unknown role for UNC-34/Ena/VASP in endodermal precursor cell apical constriction and reveal spatially distinct modes of actin assembly across the constricting cortex. Formins are enriched in the constricting apical cortex, while Arp2/3 is concentrated primarily at basolateral sites, and UNC-34 shows a region-specific functional relationship with WAVE-Arp2/3. Our findings support a model in which UNC-34 preferentially functions with WAVE-regulated branched actin networks at non-Ep-P_4_ contacts to help maintain cortical integrity. More broadly, these results show how the atlas can reveal local regulatory relationships and define how distinct actin assembly pathways are deployed across the apical cortex and cell-cell contacts, for example with the apical cortex enriched with only linear actin assembly factors.

### ERM-1 and FRM-1 maintain membrane-cortex integrity challenged by apical myosin activation in apically constricting cells

Apical constriction requires actomyosin-generated forces to be transmitted to the plasma membrane and cell-cell junctions. In *C. elegans*, the cadherin-catenin complex and SAX-7/L1CAM act redundantly at cell-cell contacts to support myosin accumulation and endodermal precursor cell internalization.^50^ We therefore asked whether proteins implicated in membrane-cortex organization also contribute to maintaining cortical integrity during apical constriction. Our localization survey identified ERM-1 and FRM-1 as candidate regulators of membrane-cortex organization that were detectable but showed distinct spatial distributions (Figure 2 and Figure 5A and 5C), prompting us to test their functions during gastrulation.

**Figure 5.**
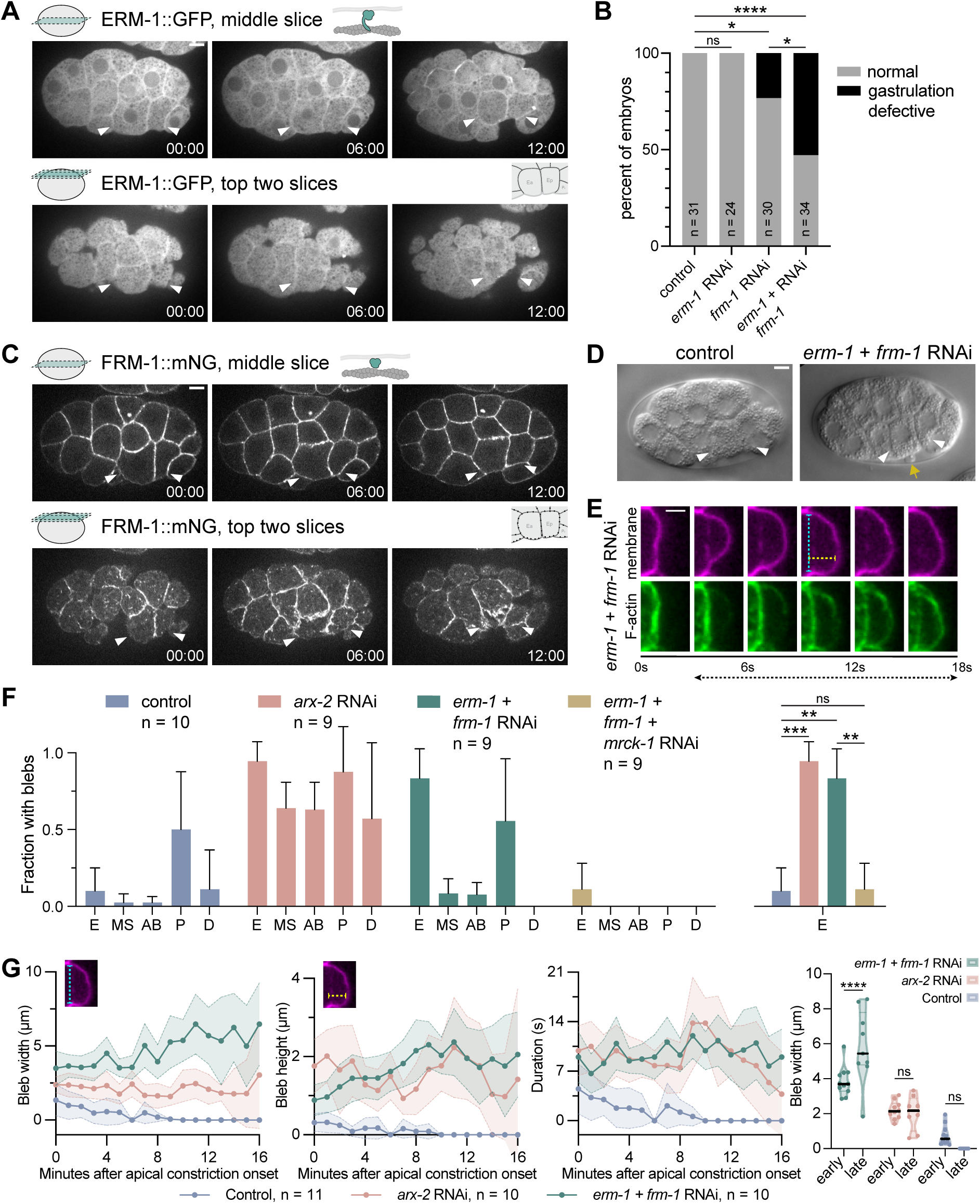
ERM-1 and FRM-1 redundantly maintain membrane-cortex integrity during apical constriction. (A) Localization of ERM-1::GFP during gastrulation. Top, optical section through the middle of the embryo; bottom, maximum-intensity projection of the two apical-most optical sections. (B) Quantification of the effects of *erm-1, frm-1,* and *erm-1 + frm-1* RNAi on gastrulation. (C) Localization of FRM-1::mNG during gastrulation. Top, optical section through the middle of the embryo; bottom, maximum-intensity projection of the two apical-most optical sections. (D) Micrographs from time-lapse DIC movies of control and *erm-1 + frm-1* RNAi embryos. Yellow arrow indicates a bleb in *erm-1 + frm-1* RNAi embryo. (E) Micrographs from a time-lapse movie of an embryo expressing mScarlet-I::PH and Utrophin::mNG following *erm-1 + frm-1* RNAi. Dashed lines illustrate the measurements used to quantify bleb morphology and dynamics: cyan, bleb width; yellow, bleb height; black with arrows, bleb duration. (F) Quantification of the fraction of cells exhibiting blebs in the E, MS, AB, P, and D lineages following the indicated RNAi treatments. (G) Left, quantification of bleb width, height, and duration following the indicated RNAi treatments. Right, comparison of bleb width between early (0–5 min) and late (11–16 min) stages following the indicated RNAi treatments. For (B), pairwise comparisons were performed using two-sided Fisher’s exact tests, with P values adjusted for multiple comparisons using the Holm–Šídák method. For (F), statistical significance was assessed using the Kruskal-Wallis test followed by Dunn’s post hoc multiple-comparisons test. For (G), changes in bleb parameters between early and late stages across RNAi conditions were analyzed using a two-way mixed-effects model, with stage (early versus late) treated as the within-embryo factor and RNAi treatment as the between-embryo factor. The full model included stage, RNAi treatment, and the stage × RNAi treatment interaction. Prespecified comparisons between early and late stages within each RNAi condition were adjusted for multiple comparisons using the Šídák method. White arrowheads indicate Ea and Ep. *P < 0.05, **P < 0.01, ***P < 0.001, ****P < 0.0001. Scale bars, 3 μm in (D) and 5 μm in all other panels. See also Figure S2 and S5.

ERM-1 is the sole *C. elegan*s ortholog of the ezrin-radixin-moesin (ERM) protein family, whose members are well-established regulators of membrane-actin cortex linkage.^51^ FRM-1 belongs to the protein 4.1/FERM-FA family and is predicted to associate with actin, although direct actin binding has not been experimentally established. Depletion of ERM-1 produced no detectable gastrulation defects, whereas depletion of FRM-1 caused mild defects (Figure 5B). Co-depletion of ERM-1 and FRM-1 significantly enhanced gastrulation defects (Figure 5B), indicating partially redundant functions during endodermal precursor cell internalization. Knockdown efficiency for each single-RNAi treatment and the combined *erm-1 + frm-1* RNAi treatment was verified using the method described above, with residual fluorescence reduced to ∼1-3% of the tagged control (Figure S3C-F). Strikingly, double-depleted embryos also formed prominent apical protrusions in Ea and Ep that resembled membrane blebs (Figure 5D, yellow arrow).

To determine whether these protrusions were bona fide blebs, we simultaneously imaged F-actin and the plasma membrane in double RNAi embryos. During protrusion formation, the membrane expanded outward with reduced cortical F-actin; after membrane expansion ceased, F-actin accumulated at the membrane, consistent with cortical reassembly during bleb maturation.^21,52^ This stereotyped sequence was observed in all embryos examined (11/11; Figure 5E), confirming that ERM-1 + FRM-1 depletion induces apical blebbing. Because blebs can result from weakened membrane-cortex attachment under cortical tension,^53^ these findings suggested that ERM-1 and FRM-1 redundantly maintain the apical membrane-cortex interface during constriction.

We and others have previously observed bleb formation in all cells after depletion of members of a Rac - WAVE - Arp2/3 pathway,^21,37,54^ prompting us to ask whether loss of the ERM-1 and FRM-1 membrane-cortex linkers produces a similarly widespread cortical defect. We found that the two perturbations produced distinct patterns of blebbing. Arp2/3 depletion increased bleb formation across multiple embryonic cell types, whereas ERM-1 and FRM-1 co-depletion caused blebbing predominantly in the apically constricting Ea and Ep cells (Figure 5F). Both ERM-1 + FRM-1 co-depletion and Arp2/3 depletion produced wider, taller, and longer-lived blebs than those seen occasionally in wild-type embryos, but their temporal patterns differed (Figure 5G). These results suggest that Arp2/3 broadly supports cortical integrity, while ERM-1 and FRM-1 are particularly important for maintaining the apical cortex in the cells undergoing apical constriction; without ERM-1 and FRM-1, apically constricting cells frequently formed wide blebs.

Our previous work showed that endodermal precursor cell internalization undergoes a mechanical transition: early actomyosin contraction generates cortical tension with limited apical surface reduction, whereas later the actomyosin network and apical cell-cell contacts move increasingly in concert.^4^ In ARX-2-depleted embryos, blebs occurred throughout internalization without a consistent change in size. In contrast, ERM-1 and FRM-1 co-depleted embryos formed progressively larger blebs, with significantly greater width and height during late (11-16 min after MSx division) vs early (0-5 min) internalization; no comparable stage-dependent increase occurred in controls or ARX-2-depleted embryos (Figure 5G and Figure S5). These findings suggest that maintaining cortical integrity becomes increasingly dependent on ERM-1 and FRM-1 as apical constriction progresses and mechanical load increases.

We tested whether blebbing in ERM-1 and FRM-1 co-depleted embryos depended on actomyosin contractility by co-depleting MRCK-1, which promotes apical myosin activation and apical cortical tension during endodermal precursor cell internalization.^38^ Knockdown efficiency for the combined *erm-1 + frm-1 + mrck-1* RNAi treatment was assessed as described above (Figure S3G–I): ERM-1 and FRM-1 fluorescence were similarly reduced to ∼2-3%, whereas MRCK-1 was reduced to ∼8%. Near-complete depletion of MRCK-1 was sufficient to abolish bleb formation in ERM-1 + FRM-1-depleted embryos (Figure 5F), indicating that bleb formation following loss of ERM-1 and FRM-1 depends on MRCK-1-dependent myosin activation.

Together, these results demonstrate that candidate regulators of membrane-cortex organization are especially important in cells undergoing apical constriction as myosin becomes activated apically, increasing cortical tension.^4,38^ ERM-1 and FRM-1 are present at similar levels in all cells (Figure 5A, C), yet in contrast to the broad requirement for Arp2/3 in suppressing blebbing in all cell types examined, ERM-1 and FRM-1 become particularly important to prevent membrane-cortical detachment in Ea and Ep as constriction progresses, dependent on the myosin activator MRCK-1. These results reveal a mechanism that enables the apical cortex to withstand additional myosin-dependent mechanical stress during apical constriction.

## Discussion

Regulation of actomyosin assembly, turnover, architecture, and force transmission underlies diverse cell movements and shape changes.^55,56^ In migrating cells, the field has developed a detailed mechanistic understanding of how local actin modulations coordinate to drive forward motion with spatial and temporal precision.^29^ In contrast, there is not yet an equally detailed view of apical constriction, despite its central role in shaping tissues and internalizing cells from the surface of embryos. A similarly detailed understanding is also lacking for epithelial morphogenesis more broadly.^57^ Here, we generated a systematic endogenous localization atlas of actin regulators during *C. elegans* gastrulation, and we used patterns emerging from the atlas to guide mechanistic experiments. Together, these analyses suggest coordinated spatial organization of actin regulators across apical, junctional, and basolateral landmarks in apically constricting cells (Figure 6). This spatial organization provides a framework for hypothesis generation. Biochemical activity defines what a protein can do, whereas localization constrains where that activity could operate. Together, these dimensions create a mechanistically informed solution space that narrows the range of plausible protein relationships and identifies testable hypotheses.

**Figure 6.**
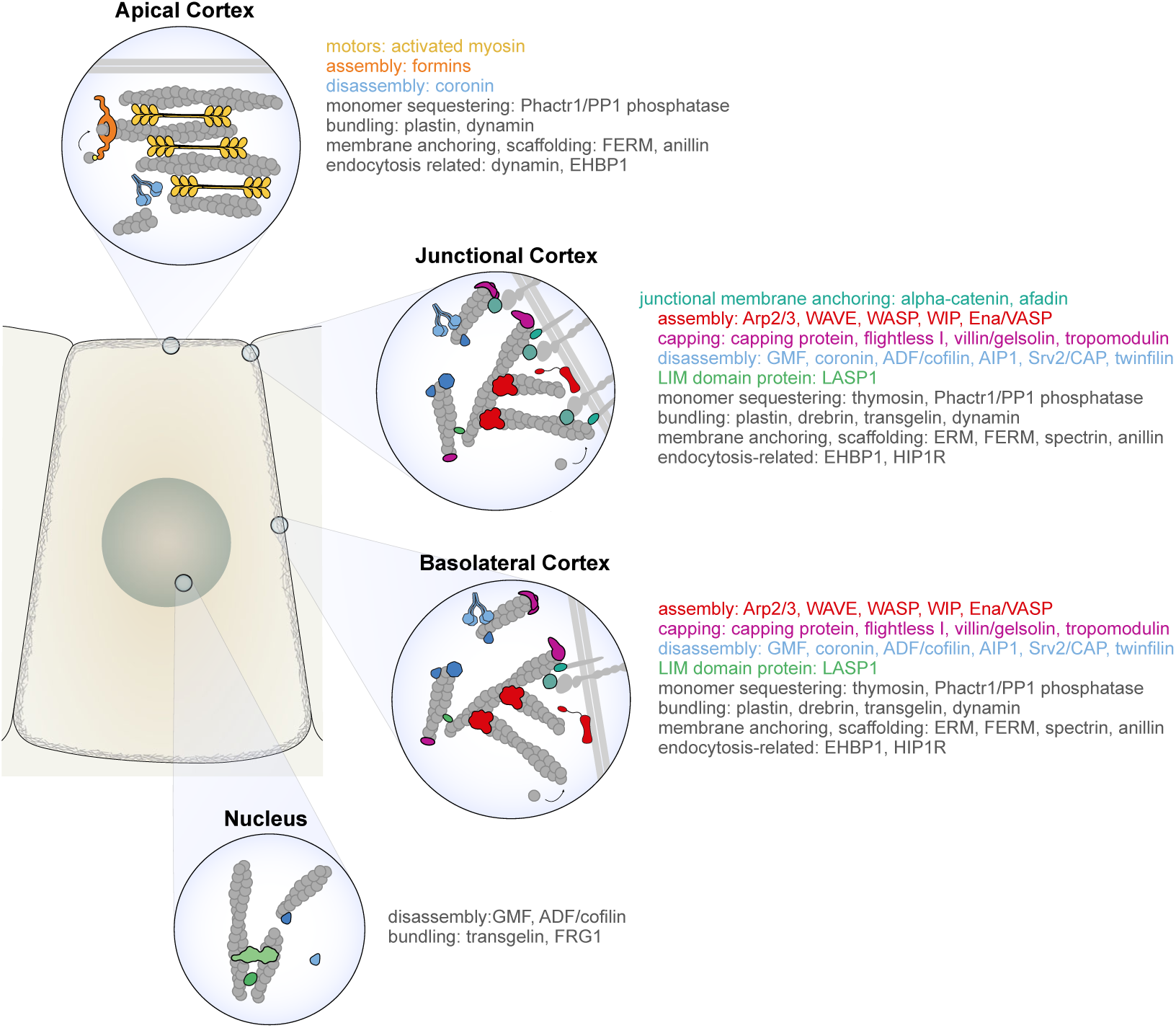
Summary: Actin regulators define landmarks in apically constricting cells. Diagram highlights some of the key differences among landmarks where ABPs were found to be enriched. ABP classes found enriched and/or functioning at specific cortical sites (apical, junctional, or basolateral) are highlighted by their inclusion in the three cortical zoom-ins and listing in colored text. Linear vs branched actin architecture diagrammed here is a working model based on which actin nucleators were found to be localized or functioning at each location. Disassemblers were found disproportionately at basolateral or junctional positions, but coronin is included apically too because there was some enrichment there above cytoplasmic level. Positions of proteins on specific filaments is only illustrative; whether capping occurs selectively on linear filaments vs branched ones is not known, for example. Activated myosin, alpha-catenin, and afadin enrichment are based on our published data; alpha-catenin and afadin are diagrammed to reflect that they become enriched junctionally during apical constriction.^24,38,65,66^ Arp2/3 is diagrammed as junctional and basolateral here based on our results here and previously,^37^ and formins are diagrammed apically because they are the sole known apical actin nucleators, although they localized to cell-cell contacts as well. Nuclear ABPs included here are the four proteins that we found enriched in nuclei above cytoplasmic level.

One such example examined here involved several ABPs whose overall distributions resembled that of the Arp2/3 component ARX-2. These proteins showed both punctate localization at cell-cell contacts and cytoplasmic puncta, suggesting that they might share an Arp2/3-dependent mechanism of spatial organization. Consistent with this prediction, representative ABPs colocalized with ARX-2, and ARX-2 depletion reduced their contact-associated enrichment and disrupted their spatial organization. Our findings reveal that basolateral cell-cell contact sites are landmarks loaded with a diversity of actin regulators, and they suggest that Arp2/3-dependent actin architecture provides a shared spatial framework for organizing regulators with distinct biochemical activities. Thus, recurrent localization patterns in the atlas can reveal candidate regulatory relationships that are not apparent from biochemical classification alone.

A second example emerged from the spatial organization of actin assembly pathways. Arp2/3 was concentrated primarily at basolateral cell-cell contacts, whereas the formins CYK-1 and INFT-2 were also enriched at the apical cortex, suggesting that distinct mechanisms of actin assembly are deployed in different cortical regions during apical constriction. Ena/VASP UNC-34 adds another layer to this organization. Ena/VASP promotes actin assembly primarily through barbed-end elongation and could function with different actin assembly pathways.^58^ The localization of UNC-34 at non-Ep-P_4_ contacts closely matched the regional activity of WAVE rather than WASP. Together with the reduction of UNC-34 enrichment after WAVE depletion, these data support a region-specific functional relationship between UNC-34 and WAVE-Arp2/3. The similar cortical blebbing caused by UNC-34 and WAVE-Arp2/3 depletion further suggests that these basolateral regulators contribute to apical constriction, at least in part, by maintaining cortical integrity.

The spatial segregation of these assembly factors may reflect critical differences in the actin architectures in each region. Formins generate linear filaments, whereas Arp2/3 generates branched networks, and these architectures differ in their interactions with myosin and their ability to support contraction.^36,59^ In reconstituted networks, formin-nucleated linear actin supports myosin-driven contraction more effectively than Arp2/3-nucleated branched actin does.^59^ This raises the possibility that apical formin activity generates an actin architecture particularly suited to constriction, whereas basolateral branched actin networks help maintain cortical integrity during cell-shape change, perhaps by reducing basolateral myosin-driven contraction and resulting internal pressure throughout the cell that can produce blebs.^37^

A third example revealed spatially and developmentally specific requirements for maintaining membrane-cortex integrity. Previous work showed that adhesion systems SAX-7/L1CAM and HMR-1/cadherin function redundantly in apical constriction.^50^ Our localization analysis identified ERM-1 and FRM-1 as candidate regulators of membrane-cortex organization with distinct spatial distributions during apical constriction. Although ERM-family proteins are well-established membrane-actin linkers, the molecular function of FRM-1 is less clear. Recent biochemical work showed that FRM-1 did not detectably bind F-actin alone but co-sedimented with F-actin in the presence of FRM-4, suggesting that FRM-1 may contribute to actin-associated organization through interactions with other proteins.^60^ Perturbation experiments revealed a strong synthetic phenotype upon co-depletion, consistent with partially redundant roles in maintaining membrane-cortex integrity. The requirement of ERM-1 and FRM-1 was strongly dependent on cellular and developmental context. Loss of ERM-1 and FRM-1 preferentially destabilized the cortex in the constricting endodermal precursor cells, and the defect became more pronounced as internalization progressed. During this period, actomyosin contractions and movement of apical cell-cell contacts become increasingly coordinated.^4^ Together with the suppression of blebbing by reduced apical myosin activation, these findings suggest that ERM-1-and FRM-1-dependent membrane-cortex coupling becomes increasingly important as contractile activity and movement of apical cell contacts become more closely coordinated. In contrast, the broader blebbing caused by Arp2/3 depletion indicates that cortical integrity is maintained by multiple spatially and mechanistically distinct systems. These results suggest that ERM-1 and FRM-1 maintain membrane-cortex connections that are particularly challenged by apical myosin-dependent forces during apical constriction.

Beyond the mechanistic insights generated in this study, this work establishes a resource for the field: a collection of sequence-verified, endogenously tagged ABPs that were systematically evaluated for evidence of tag-induced functional disruption, and a visual imaging atlas documenting ABP localization in the early embryo. By interpreting localization patterns as a solution space that reveals emergent functional hubs of actin regulators, and by combining these observations with functional perturbation, we outline a systems-cell-biology framework for mechanistic insight into apical constriction using the *C. elegans* embryo. This approach may be particularly useful for proteins whose essential functions or redundancy complicate conventional loss-of-function studies, especially when combined with perturbations that provide greater temporal or cell-type specificity.^61–64^ This approach is broadly applicable to other tissues and provides a foundation for dissecting actin-dependent processes using *C. elegans* as a versatile experimental platform.

## Materials and methods

### Strain maintenance

Table S2 summarizes the *C. elegans* strains used in this study and their sources. Worms were maintained on Nematode Growth Medium (NGM) plates at 20-22 °C using standard culturing methods.^67^

### CRISPR-mediated genome engineering

Endogenously tagged strains were generated using a previously described CRISPR/Cas9-mediated genome editing protocol.^68^ Cas9 target sequences were designed with CRISPOR ^69^. tracrRNA and crRNA were combined with Cas9 protein (IDT) and incubated at 37°C for 15 min to assemble ribonucleoprotein complexes. Donor DNA containing codon-optimized mNeonGreen or mScarlet-I (amplified from pDD268 and pMS050, respectively) with 35 bp homology arms was denatured, cooled, and added to the mixture together with the co-injection marker plasmid pRF4 [*rol-6(su1006)*]. The injection mix was centrifuged, transferred to a fresh tube, and kept on ice until use. Heterozygous F1 progeny were identified by genotyping or fluorescence screening using a Zeiss AxioZoom V16 stereo microscope, and homozygous lines were sequence-verified. PHX strains were generated by SunyBiotech.

### RNA interference

cDNA from wild-type N2 worms was synthesized using SuperScript III First-Strand Synthesis SuperMix (Invitrogen). Each primer included 15 bases of the T7 promoter at the 5′ end for a two-step PCR using wild-type cDNA as the template. PCR products were purified with the Zymoclean Gel DNA Recovery Kit (Zymo Research) and used as templates for a second PCR with primers containing the full-length T7 promoter sequence. The purified PCR products were then used as templates for in vitro transcription with the T7 RiboMAX Express RNAi System (Promega) following the manufacturer’s instructions.

Purified dsRNA (300–1,000 ng/μL) was injected into L4 animals using a Narishige injection apparatus, a Parker Instruments Picospritzer II, and a Nikon Eclipse TE300 microscope with DIC optics. Aliquots of dsRNA were stored at-80 °C to avoid repeated freeze-thaw cycles. Injected worms were recovered on seeded NGM plates for at least 28 h at room temperature before embryos were harvested for imaging.

### Image acquisition

Embryos were dissected and mounted on poly-L-lysine-coated coverslips in *C. elegans*egg buffer on 2% agar pads, then sealed with VALAP (1:1:1 vaseline, lanolin, and paraffin). Fluorescence imaging was performed on Nikon Eclipse Ti inverted confocal microscopes equipped with Yokogawa CSU-X1 spinning-disk scan heads and either a Hamamatsu ORCA-Quest qCMOS camera or a Hamamatsu ORCA-Flash4.0 sCMOS camera. Images were acquired at room temperature using a 60× 1.4 NA oil-immersion objective with 488-, 514-, and 561-nm lasers; some images were taken with a 1.5× magnifier. Z-stacks were collected with a 0.5-1 µm step size. Four-dimensional DIC imaging was performed on a Nikon Eclipse 800 microscope with a Diagnostic Instruments SPOT2 camera, capturing 1-µm optical sections every minute using a 60× 1.4 NA oil objective. Near-TIRF imaging of CYK-1::GFP was performed on a Nikon Ti2-E inverted microscope as previously described.^70^ Image acquisition was carried out using MetaMorph or NIS-Elements software.

### Colocalization analysis

Colocalization of proteins was quantified in Fiji/ImageJ^71^ using the Coloc 2 plugin. Embryos were analyzed 6 min after completion of MSx division. For each region analyzed, a single z-plane showing the clearest localization of the proteins of interest was selected. Each channel was processed independently by subtracting the background using a rolling-ball radius of 50 pixels, followed by Gaussian blurring with a radius of 2 pixels. Colocalization was measured within defined regions of interest (ROIs). For cell-cell contacts, a 12-pixel-wide freehand line was traced along either the Ep-P_4_ contact or other cell-cell contacts. When multiple non-Ep-P_4_ contacts were analyzed together, individual ROIs were combined into a single ROI. For cytoplasmic measurements, freehand ROIs encompassing the cytoplasm of Ea and Ep were drawn while excluding the nuclei and then combined into a single ROI. The processed image pairs were analyzed with Coloc 2 using the selected ROI and Costes threshold regression. For negative controls, the image for a single channel was rotated 180°, and colocalization was measured using the same ROI as for non-Ep-P_4_ contacts. Pearson’s correlation coefficient (PCC) calculated without intensity thresholding was used as the measure of colocalization.

### Quantitative measurements of protein localization along cell-cell contacts

Line-scan analysis was performed 6 min after completion of MSx division. For each embryo, a z-plane containing prominent puncta along the Ea-Ep contact was selected. A 3-pixel-wide line was traced along the Ea-Ep interface in Fiji/ImageJ, and fluorescence intensity profiles along the same line were collected from the ARX-2 and the other channel using Plot Profile. Mean cytoplasmic fluorescence was measured using a 3-pixel-wide line positioned away from bright puncta or other structures, and background fluorescence was measured from an off-embryo region using the same line width. Cytoplasmic and background measurements were collected for both fluorescence channels and used in the subsequent processing of the intensity profiles.

### Quantification of RNAi efficiency

To assess RNAi knockdown efficiency, stage-matched wild-type, knock-in control, and RNAi-treated embryos were mounted and imaged together under the same conditions. For each embryo, a sum-intensity projection of the fluorescence z-stack was generated in Fiji/ImageJ. Mean fluorescence intensity was measured within an ROI outlining the embryo, and background intensity was measured from an area outside the embryo in the same image. Background-corrected fluorescence was calculated by subtracting the background value from the embryo intensity and was normalized to the corresponding control to determine the percentage of residual fluorescence.

### Quantification of changes in protein localization following RNAi treatment

Protein localization was quantified 6 min after completion of MSx division. For each region analyzed, a single z-plane that best resolved the relevant localization was selected in Fiji/ImageJ. Cell-cell contacts were traced using a 3-pixel-wide freehand line, with the Ep-P_4_ contact and other cell-cell contacts analyzed separately. Cytoplasmic ROIs were drawn within Ea and Ep while excluding the nuclei, and the two ROIs were combined for analysis. Additional ROIs were drawn around the entire embryo and in an off-embryo background region.

Mean fluorescence intensity was measured for each ROI. For each embryo, the mean intensity of the background ROI was subtracted from all other measurements. Regional intensities were then normalized to the whole-embryo fluorescence of each embryo.

### Quantification of the cellular distribution of bleb formation

To assess the cellular distribution of bleb formation, z-stacks were acquired at 1-min intervals with a 0.5 µm step size. At selected time points, Ea, Ep, and all other cells with exposed cortical surfaces were examined throughout the z-stack for bleb formation. A cell was scored as bleb-positive when a clear separation between the plasma membrane and the underlying cortex was observed. For each cell type, the fraction of cells exhibiting a bleb was calculated.

### Quantification of bleb width, height, and duration

Bleb width, height, and duration were quantified from time-lapse movies acquired at 3 second intervals. The timing of MSx, ABx, and E cell divisions was recorded when visible. Movies were divided into consecutive 20-frame (1-min) intervals, and the largest bleb formed on Ea within each interval was identified. Bleb duration was measured from the frame in which the plasma membrane first detached from the underlying actin cortex to the frame in which cortical actin re-established contact with the membrane. The maximum width and height reached by each bleb before reattachment were measured independently and could be obtained from different frames. Intervals in which no bleb was observed were assigned a width, height, and duration of zero. Measurements were aligned relative to AB cell division, which provided a consistent temporal reference across movies.

### Statistical analysis

Statistical analyses were performed using GraphPad Prism 10. For continuous numerical data, statistical tests were selected based on residual normality and equality of variance. Residual normality was assessed using the Shapiro-Wilk test for both two-group comparisons and one-way ANOVA. Equality of variance was evaluated using an F test for two-group comparisons and a Brown-Forsythe test for one-way ANOVA.

For comparisons between two groups, an unpaired t-test was used when residuals were normally distributed and variances were comparable. Welch’s t-test was used when residuals were normally distributed but variances differed, whereas the Mann-Whitney test was used when residuals deviated significantly from normality.

For comparisons among multiple independent groups, ordinary one-way ANOVA followed by Tukey’s multiple-comparisons test was used when residuals were normally distributed and variances were comparable. Brown-Forsythe and Welch ANOVA was used when residuals were normally distributed but variances differed, and the Kruskal-Wallis test was used when residuals deviated significantly from normality.

For colocalization analyses, Pearson correlation coefficients (PCCs) were Fisher r-to-z transformed before statistical testing, while untransformed PCCs were displayed in the figures. When PCCs from multiple cellular regions were measured within the same embryo, Fisher-transformed values were analyzed using repeated-measures one-way ANOVA with Geisser-Greenhouse correction, followed by Dunnett’s multiple-comparisons test against the prespecified reference group. The distribution of residuals from repeated-measures analyses was evaluated using quantile-quantile plots. Comparisons of Fisher-transformed PCCs among independent protein groups were performed using ordinary one-way ANOVA followed by Dunnett’s multiple-comparisons test against the prespecified reference group.

Changes in bleb parameters between early and late stages across RNAi conditions were analyzed using a two-way mixed-effects model, with stage (early versus late) treated as the within-embryo factor and RNAi treatment as the between-embryo factor. The full model included stage, RNAi treatment, and the stage × RNAi treatment interaction. Prespecified comparisons between early and late stages within each RNAi condition were adjusted for multiple comparisons using the Šídák method. Statistical significance was assessed at α = 0.05.

Categorical data were analyzed using two-sided Fisher’s exact tests. For experiments involving multiple pairwise comparisons, P-values were adjusted within each prespecified family using the Holm-Šídák method, with the family-wise error rate set at 0.05. All other statistical comparisons were also two-sided, and the tests used for individual experiments are indicated in the corresponding figure legends.

## Data visualization

Representative frames from time-lapse movies were processed in ImageJ/Fiji. Figures were assembled in Adobe Illustrator and GraphPad Prism 10.

## Acknowledgments

We thank Dan Dickinson, David Kovar, Katherine Long, Amy Maddox, and Ed Munro for discussions and comments on the manuscript. We thank Yucheng Zhang for assistance. This work was supported by NIH grant R35GM134838 to B.G., NIH T32GM133364 (B.N.d.J.), Chan Zuckerberg Initiative (D.Q.M.), Human Frontier Long Term Fellowship LT000127/2016 (C.Y.), NIH R35GM142880 (A.M.P.), NIH F31HD112152 (T.V.G.), and American Cancer Society postdoctoral fellowship A24-0591-001 and the L’Oréal For Women in Science Program (T.N.M.-K.). Some strains were provided by the CGC, which is funded by NIH Office of Research Infrastructure Programs (P40 OD010440).

## Supplementary figure and table legends

**Figure S1.**
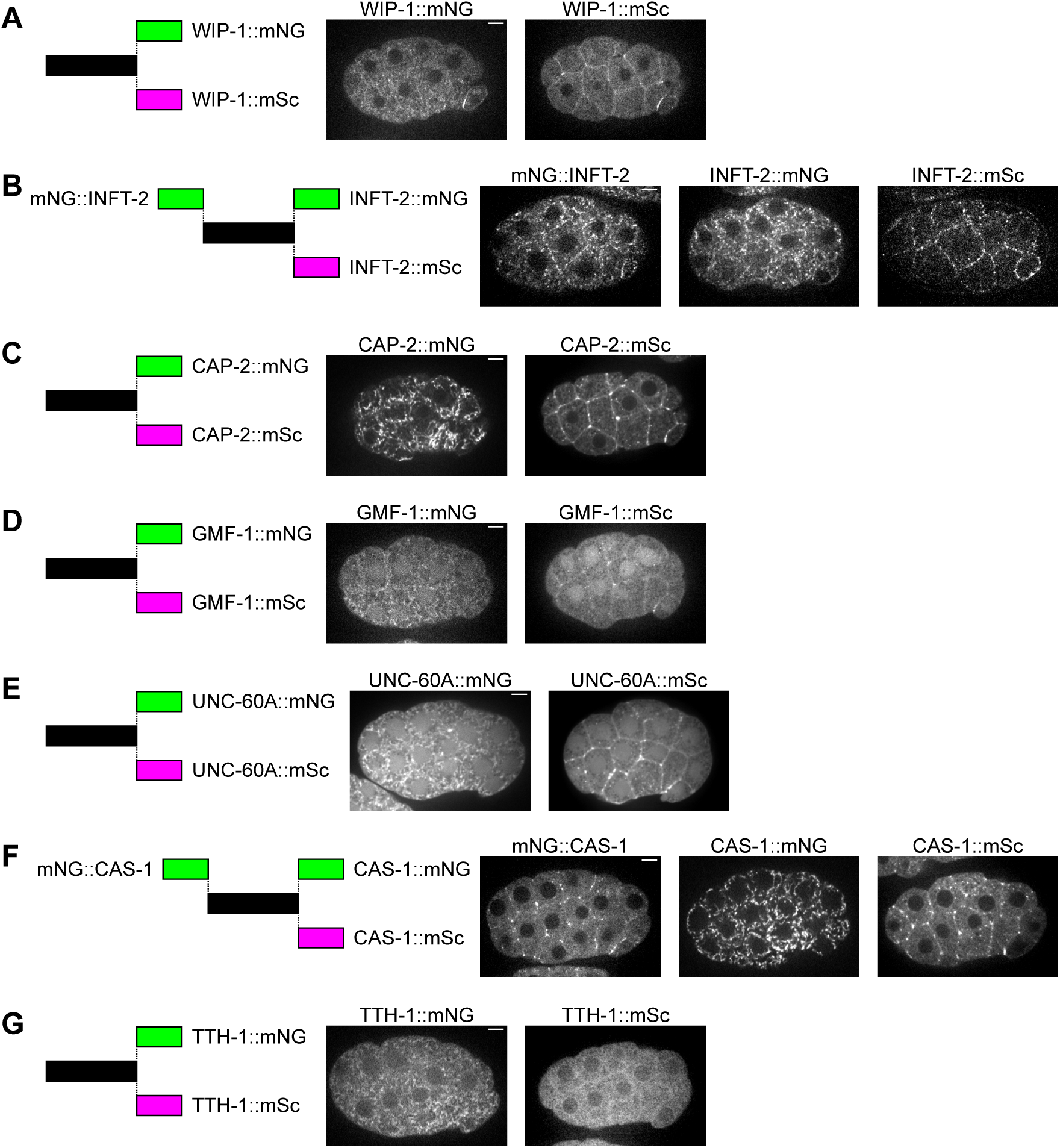
Changing fluorescent-tag configurations resolves unexpected localization patterns, related to. Figure 2. (A–G) Comparison of localization patterns among different fluorescent tag configurations. For each protein, diagrams on the left indicate the fluorescent protein and tag position for each endogenously tagged allele, and images on the right show the corresponding localization. Scale bars, 5 µm.

**Figure S2.**
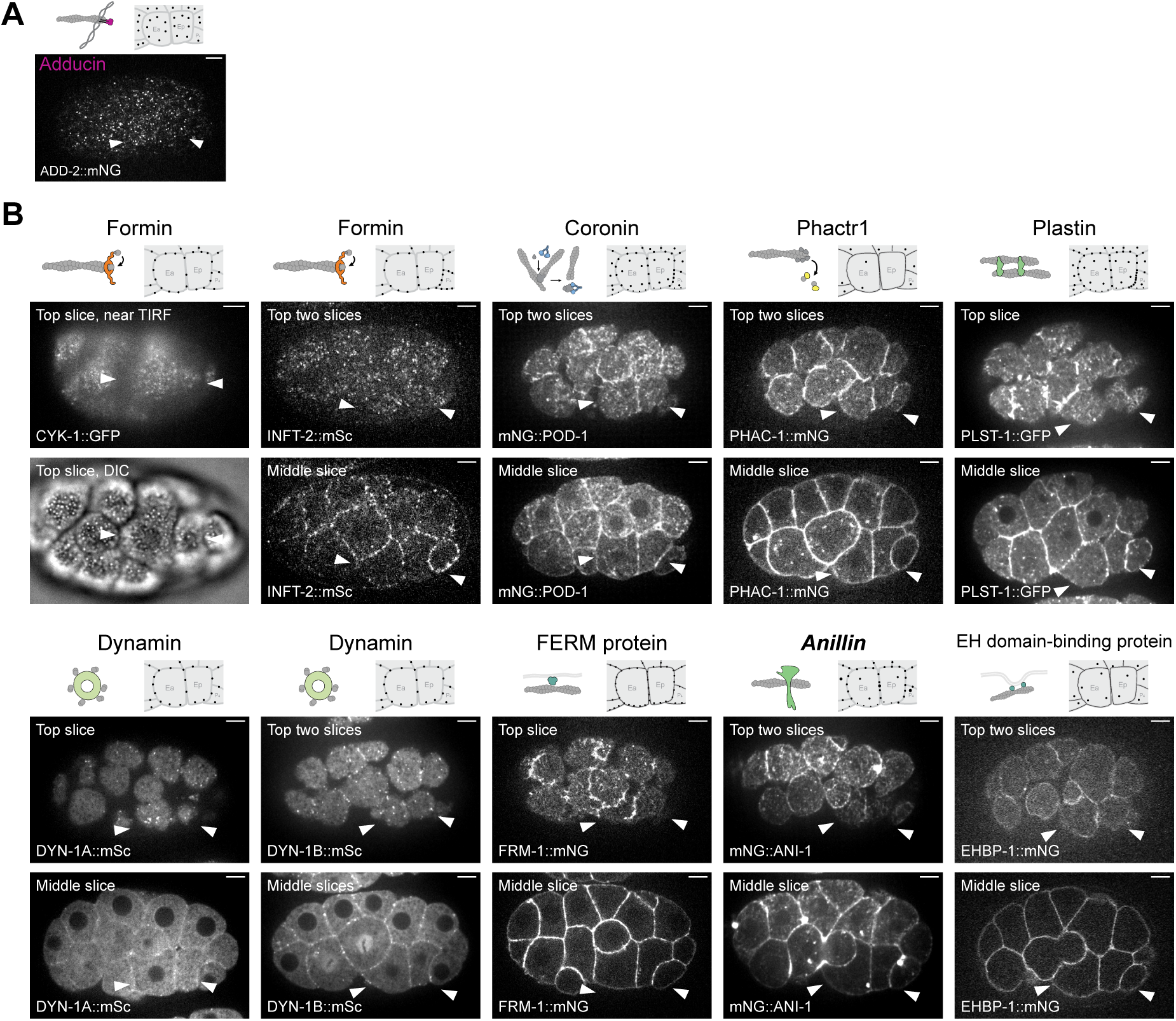
Localization of ADD-2 and distribution of selected ABPs at the cell apex, related to. Figure 2 **and 4**. (A) Localization pattern of ADD-2::mNG. Upper left, diagram depicting reported interactions with F-actin; upper right, schematic summarizing the major localization patterns in Ea, Ep, and neighboring cells; bottom, maximum-intensity projection of an embryo expressing endogenously tagged ADD-2::mNG. (B) Apical localization of ABPs. Upper left, diagrams depicting reported interactions with F-actin; upper right, schematics summarizing the major localization patterns in Ea, Ep, and neighboring cells; bottom, images of embryos expressing endogenously tagged proteins. For CYK-1, the top row shows the apical-most optical section, and the bottom row shows the corresponding DIC image. For the other proteins, the top row shows either the apical-most optical section or a maximum-intensity projection of the two apical-most sections, and the bottom row shows a middle optical section. Near TIRF is used for CYK-1 to show apical distribution with high signal/background ratio. White arrowheads indicate Ea and Ep. Scale bars, 5 µm.

**Figure S3.**
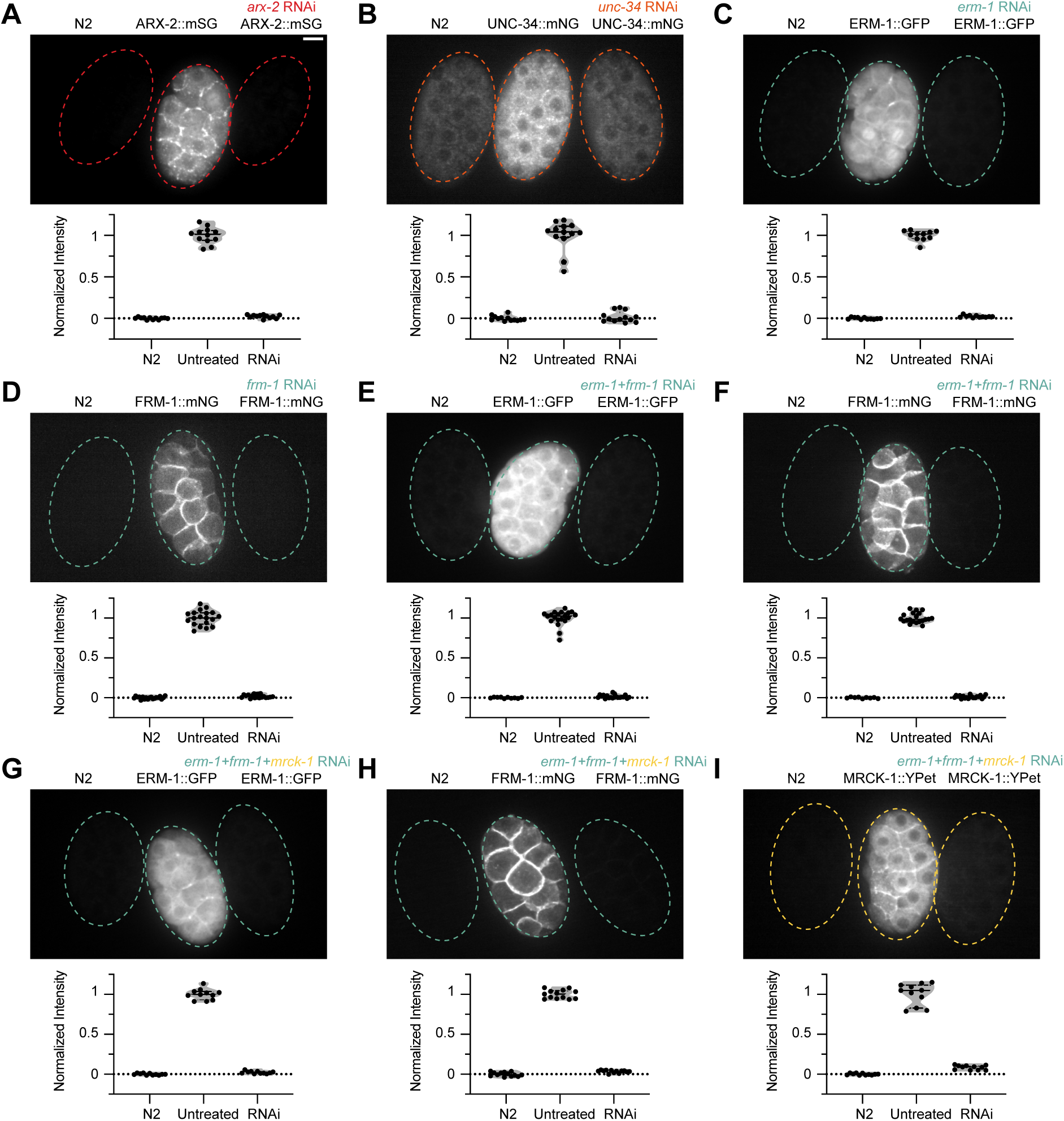
Validation of RNAi-mediated depletion of ABPs, related to. Figure 3**, 4, and 5.** (A-I) Top, fluorescence micrographs of stage-matched wild-type, knock-in, and RNAi-treated embryos mounted side-by-side and imaged from a lateral view. Bottom, violin plots showing normalized whole-embryo fluorescence intensity in wild-type, knock-in, and RNAi-treated embryos. Mean fluorescence intensity was normalized such that the average intensity of wild-type embryos was set to 0 and that of knock-in embryos was set to 1. Scale bars, 5 µm.

**Figure S4.**
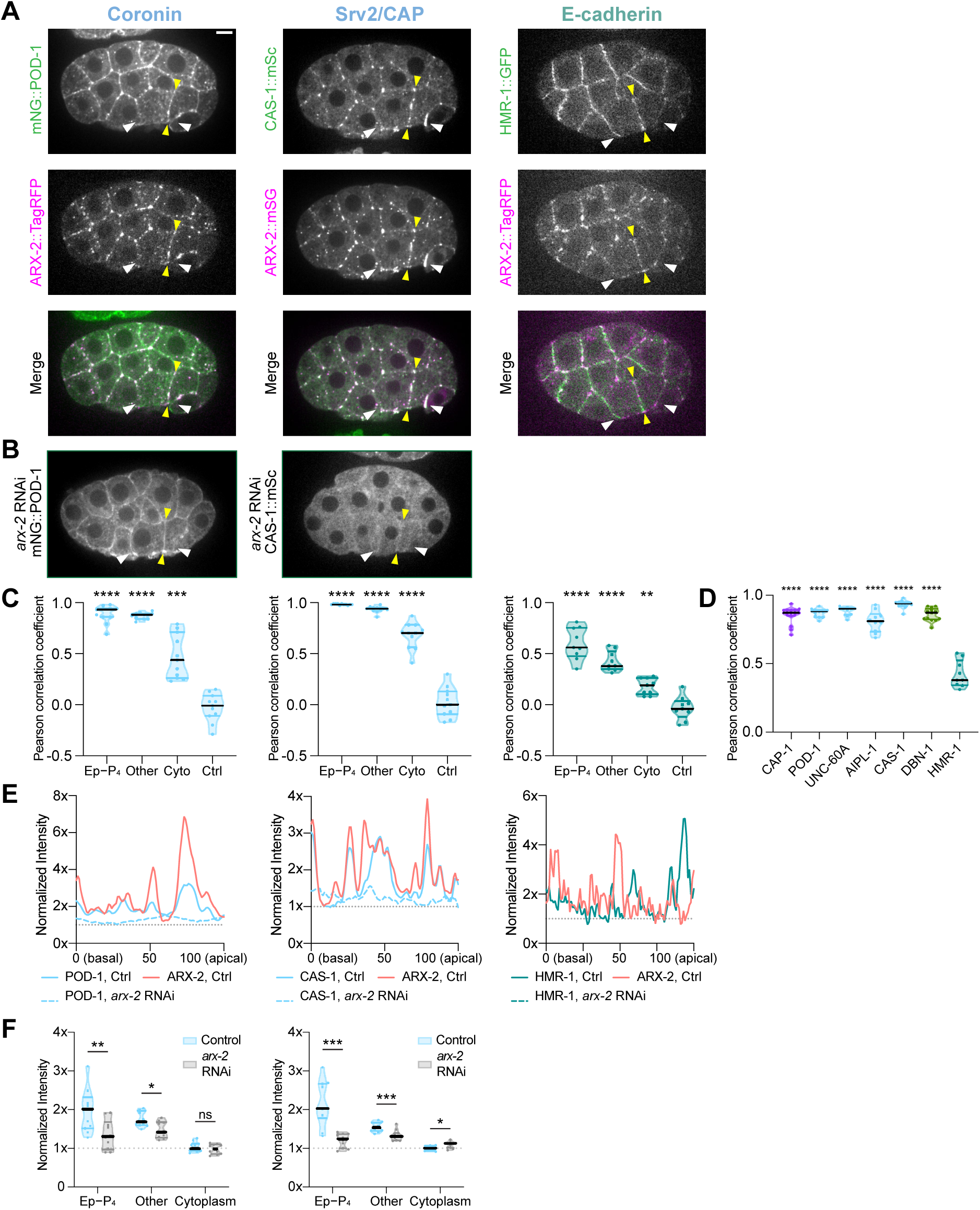
Additional analysis of actin regulators at cell-cell contacts, related to. Figure 3. (A) Co-visualization of mNG::POD-1, CAS-1::mSc, and HMR-1::GFP with ARX-2::TagRFP/mSG. (B) Localization of mNG::POD-1 and CAS-1::mSc following *arx-2* RNAi. (C) Quantification of colocalization between each protein of interest and ARX-2. Violin plots show PCCs measured at the Ep-P_4_ contact, other cell-cell contacts, the cytoplasm, and a negative control generated by rotating one channel 180°. PCCs were Fisher *r*-to-*z* transformed before statistical analysis. (D) Comparison of PCCs at other cell-cell contacts from Figures 3C and S3C between selected ABPs and HMR-1. PCCs were Fisher *r*-to-*z* transformed before statistical analysis. (E) Representative intensity profiles along the Ea-Ep contact. Colors indicate proteins; solid and dashed lines indicate control and *arx-2* RNAi conditions, respectively. (F) Quantification of POD-1 and CAS-1 localization at the Ep-P_4_ contact, other cell-cell contacts, and the cytoplasm in control and *arx-2* RNAi-treated embryos. For (C), Fisher-transformed PCCs were analyzed using repeated-measures one-way ANOVA with Geisser-Greenhouse correction followed by Dunnett’s multiple-comparisons test comparing each cellular region with the negative control. For (D), Fisher-transformed PCCs were analyzed using one-way ANOVA followed by Dunnett’s multiple-comparisons test comparing each selected actin regulator with HMR-1. For (F), comparisons between control and *arx-2* RNAi were performed using an unpaired t test, an unpaired t test with Welch’s correction, or a Mann-Whitney test, as appropriate based on data distribution and variance. White arrowheads indicate Ea and Ep; yellow arrowheads indicate the Ea-Ep contact. *P < 0.05, **P < 0.01, ***P < 0.001, ****P < 0.0001. Scale bars, 5 μm.

**Figure S5.**
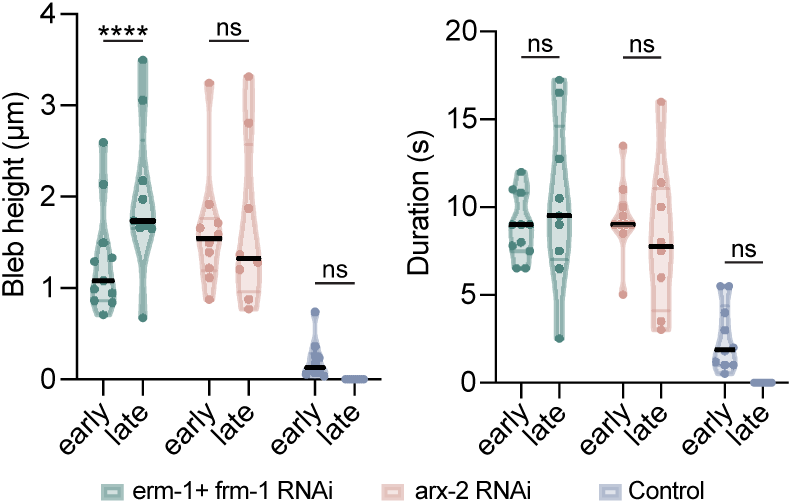
Stage-dependent changes in bleb height and duration following RNAi treatments, related to. Figure 5. Comparison of bleb height and duration between early (0-5 min) and late (11-16 min) stages following the indicated RNAi treatments. Changes in bleb height and duration between early and late stages across RNAi conditions were analyzed using a two-way mixed-effects model, with stage (early versus late) as the within-embryo factor and RNAi treatment as the between-embryo factor. The model included stage, RNAi treatment, and the stage × RNAi treatment interaction. Prespecified comparisons between early and late stages within each RNAi condition were adjusted for multiple comparisons using the Šídák method. ****P < 0.0001.

---

**Table S1.**
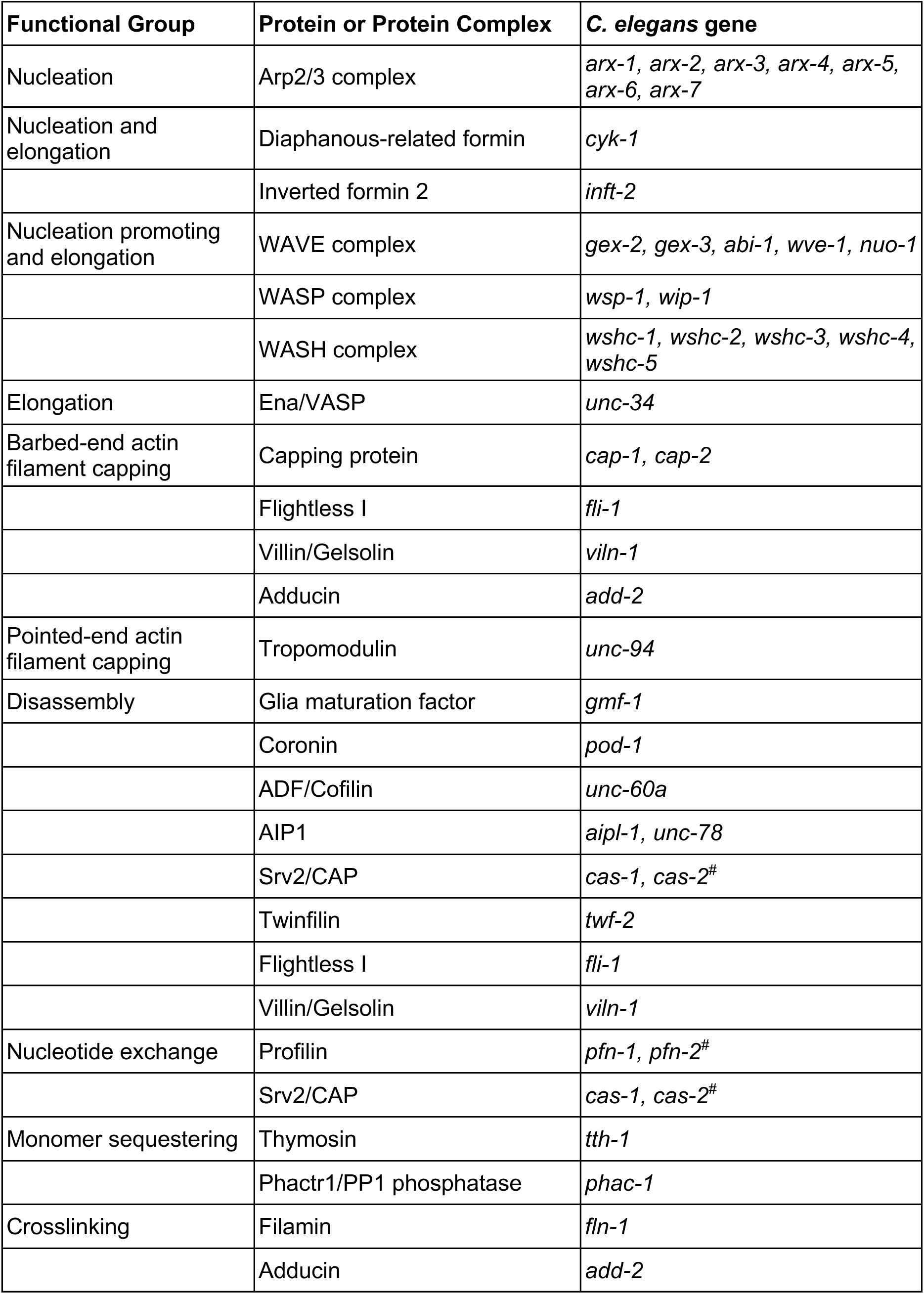
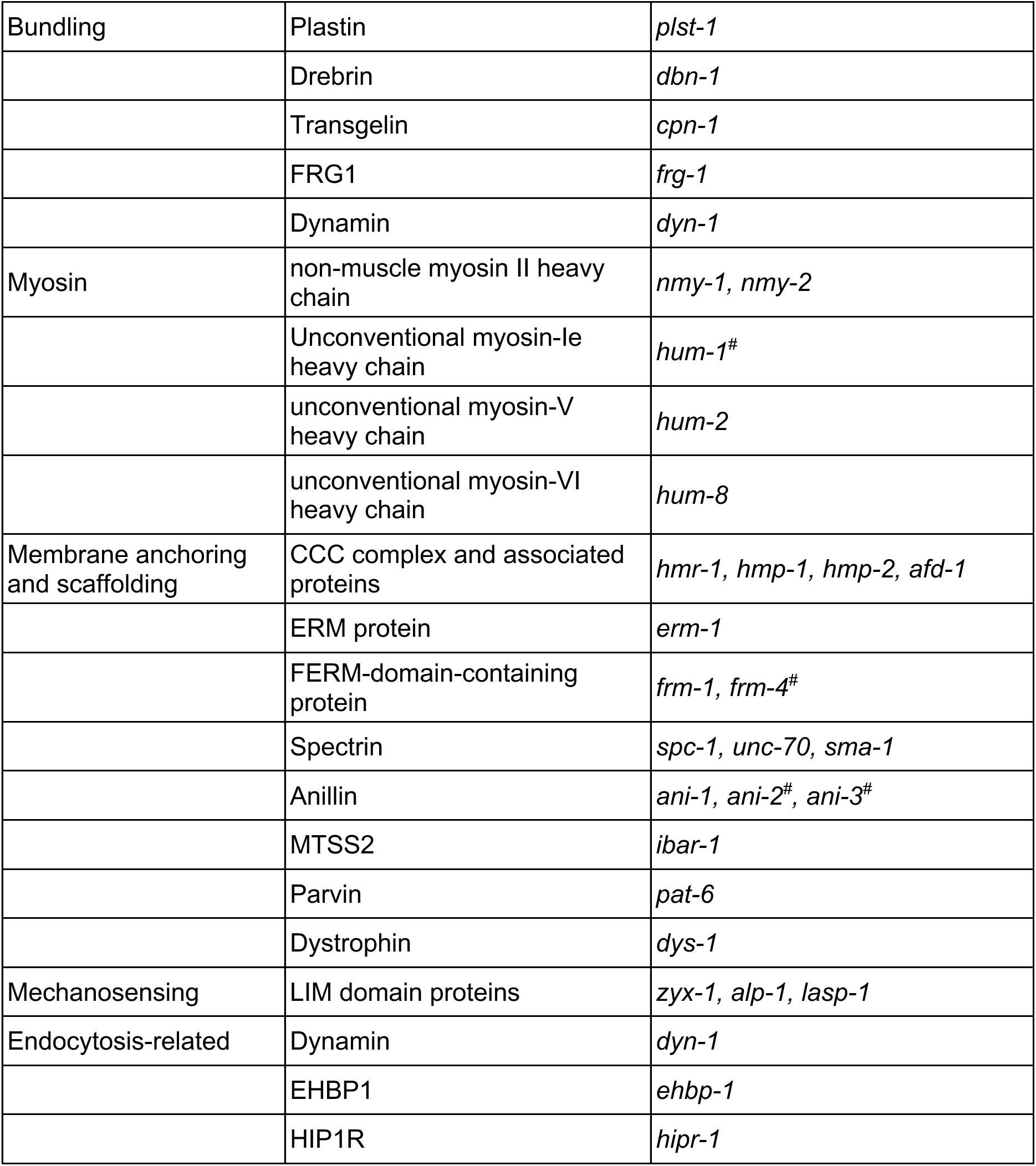
Functional classes of *C. elegans* ABPs candidates, related to Figure 1. ABP candidates included in Table S1 are organized according to their known or proposed molecular functions in actin regulation. Proteins not characterized either because another family member was selected or due to technical limitations, are marked with #. Additional proteins identified through actin-binding GO annotations but lacking well-established actin-regulatory functions in *C. elegans* (F54F2.7, MAPH-1.2, INX-3, IPLA-2/3, T24F1.2, HCP-2, and VPS-16) were not included in Table S1.

**Table S2.**
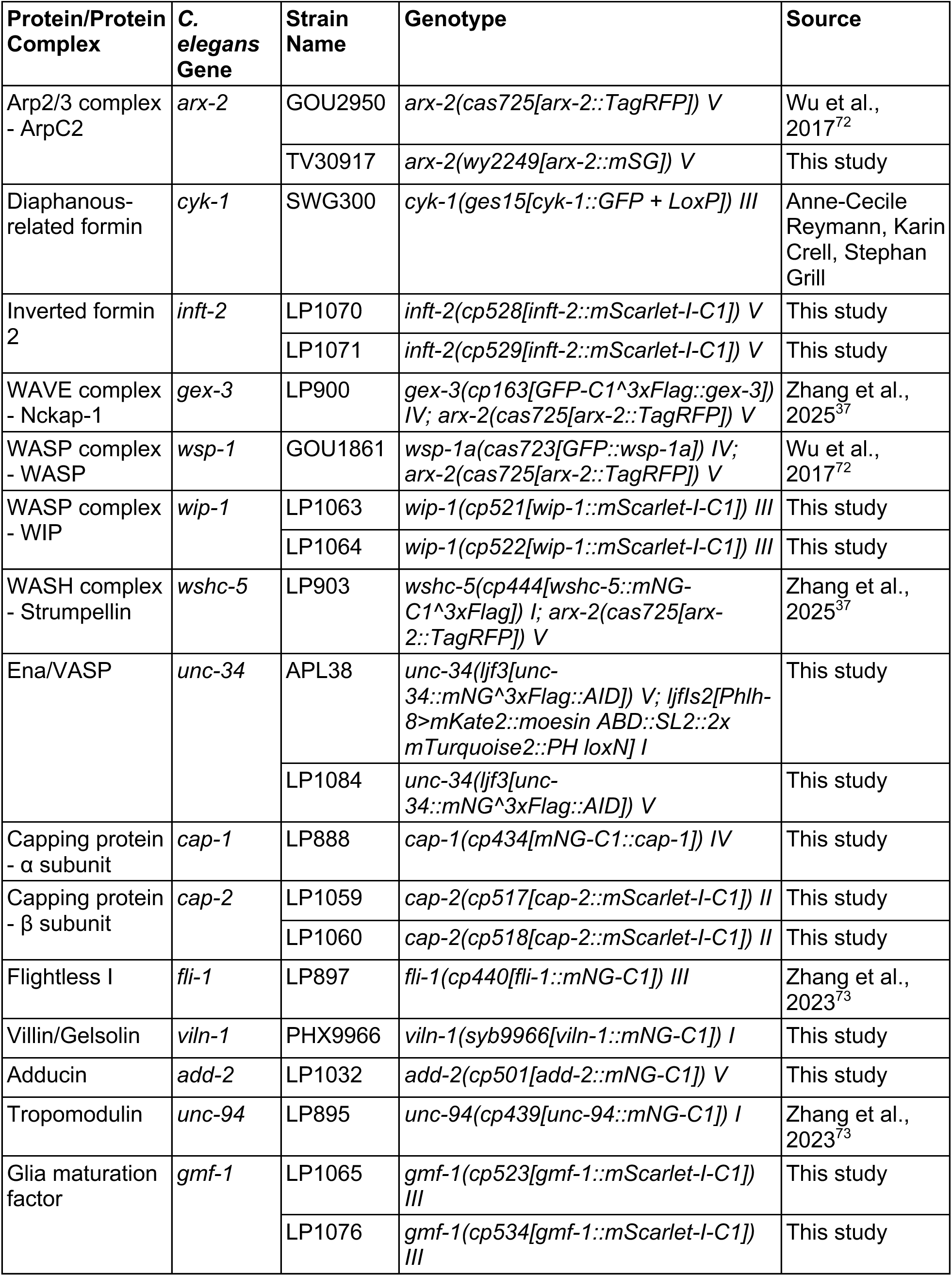
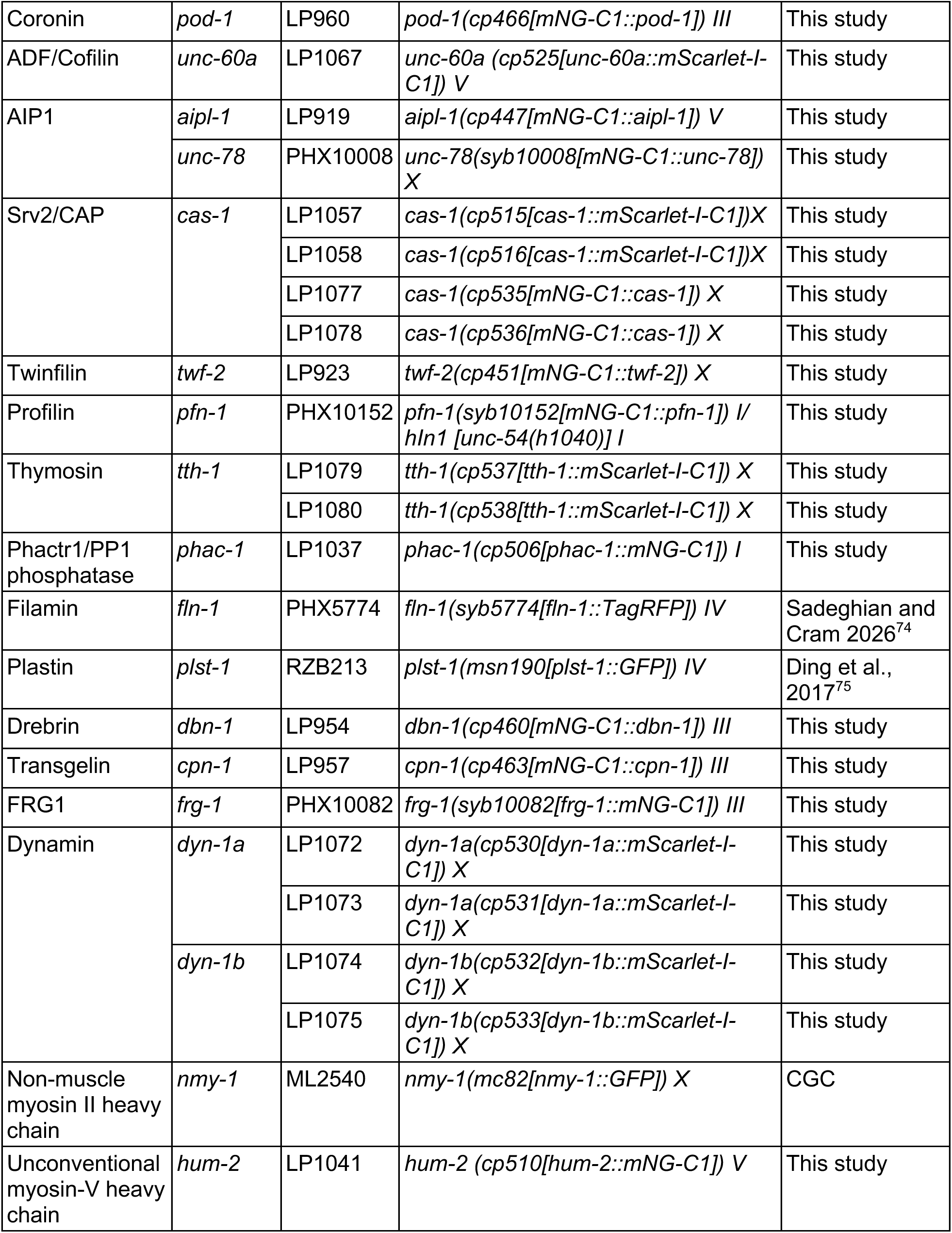
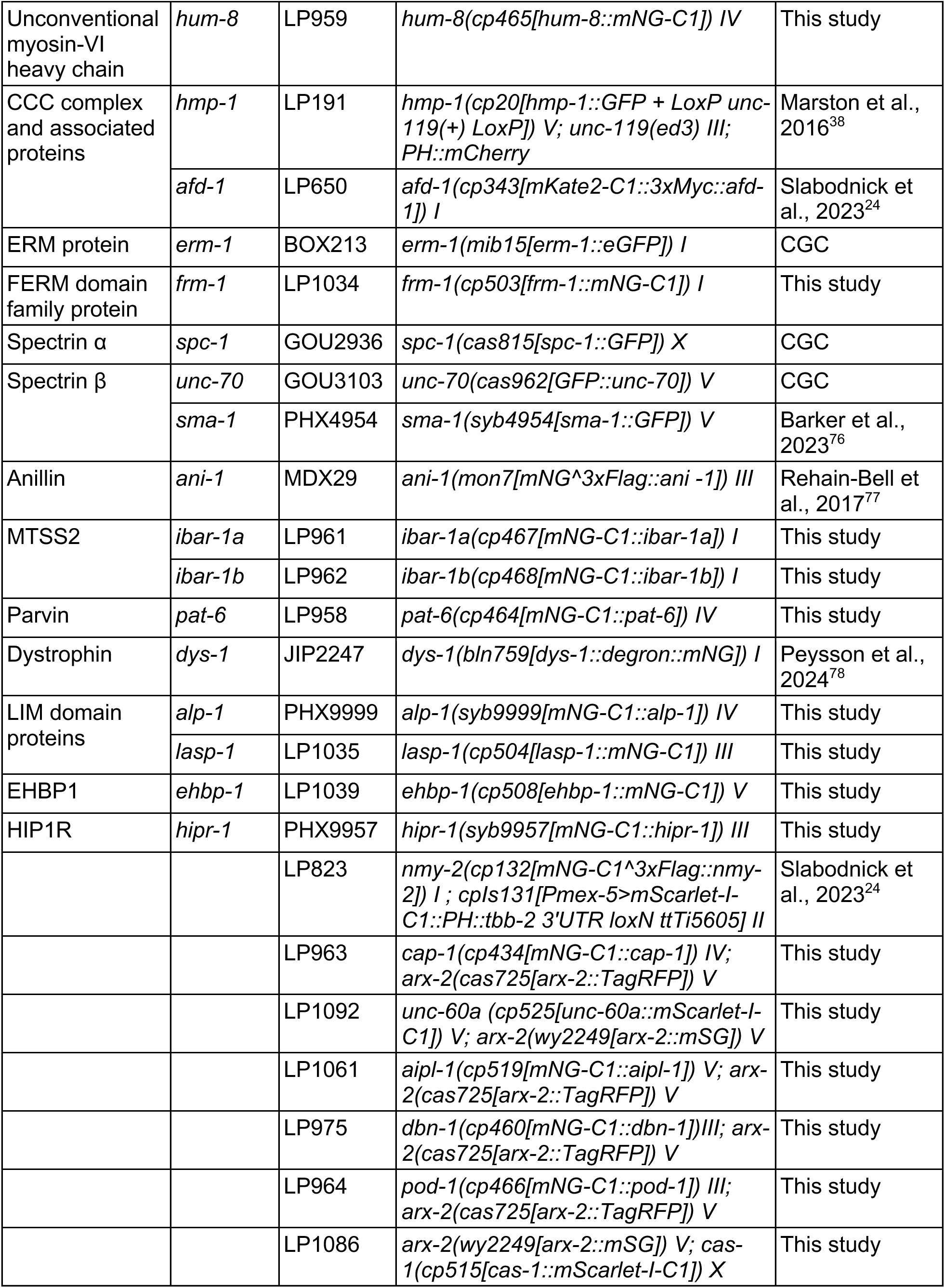
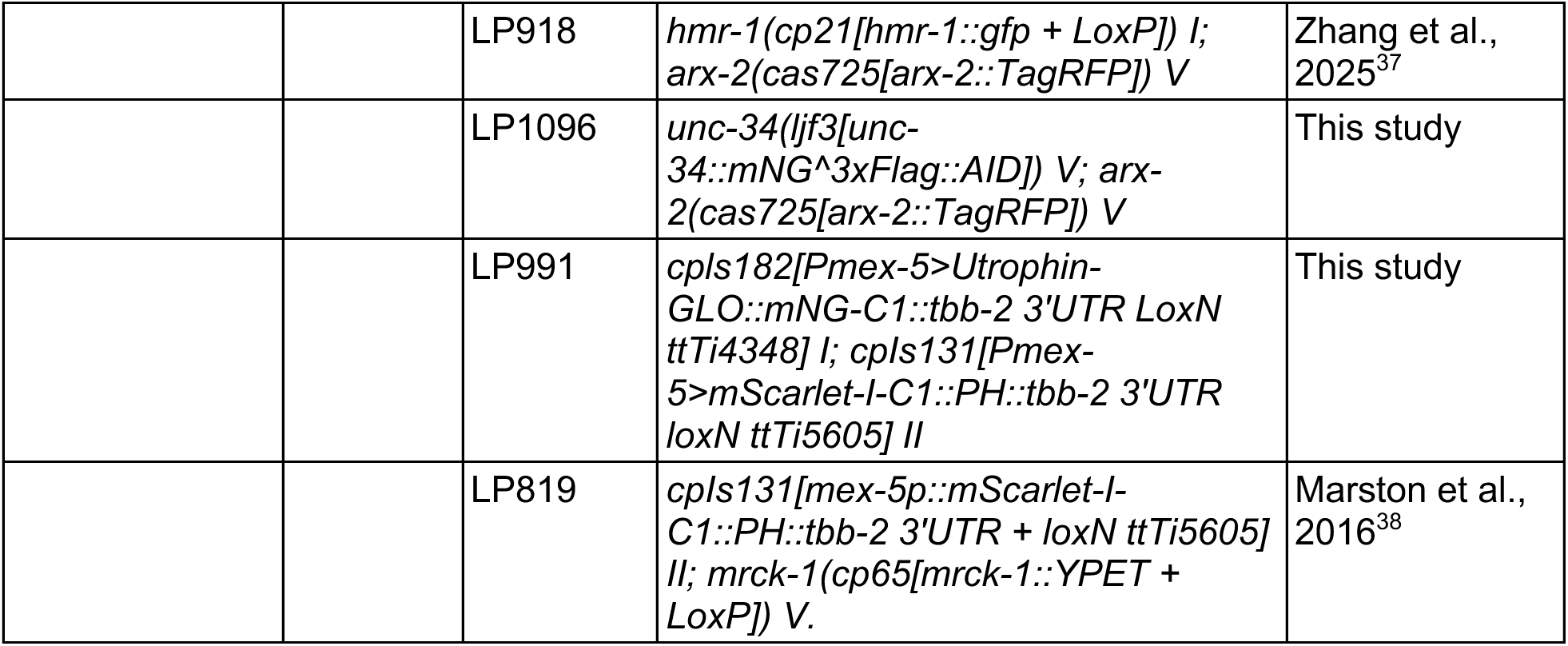
Strains, related to Figure 2 and Table 1. Names, genotypes, and sources for all strains used in this study.

**Table S3.** Evaluation of tagged ABPs, related to Figure 2 and Table 1. Summary of hatching rates, known localization, and known loss-of-function phenotype associated with each ABP.

| ***C. elegans* Gene** | **Strain Name** | **Hatching Rate** | **Previously reported localization** | **Known Phenotype** |
| --- | --- | --- | --- | --- |
| *arx-2* | GOU2950 | 100% | Localizes to cell-cell contacts as well as vesicle-like structures in the cytoplasm in *C. elegans* embryos<sup>37</sup> | RNAi: >95% embryonic lethal<sup>20</sup> |
|  | TV30917 | 99% |  |  |
| *cyk-1* | SWG300 | 100% | Localizes to cortex with enrichment at cleavage furrow in *C. elegans* zygotes<sup>30,79</sup> | *cyk-1(or36)* embryos fail first mitosis and fail to extrude polar bodies<sup>30</sup> |
| *inft-2* | LP1070 | 99% | Transgenic INFT-2::GFP and Venus::INFT-2 colocalize with actin at the apical membrane and leading edge of *C. elegans* excretory cell<sup>80</sup> | No known embryonic phenotype |
|  | LP1071 | 100% |  |  |
| *gex-3* | LP431 | 99% | Localizes to cell-cell contacts, except at the Ep-P<sub>4</sub> contact in *C. elegans* embryos<sup>37,49</sup> | RNAi: 100% embryonic lethal<sup>81</sup> |
| *wsp-1* | GOU2049 | 100% | Localizes to cell-cell contacts, with the strongest enrichment at the Ep-P<sub>4</sub> contact in *C. elegans* embryos<sup>37</sup> | RNAi: 35% embryonic lethal<sup>82</sup> |
| *wip-1* | LP1063 | 99% | Immunostaining shows expression in all cells, primarily at cell-cell contacts, in embryos undergoing ventral enclosure<sup>83</sup><br>Co-localization with basolateral DBN-1 and DYN-1 puncta in *C. elegans* intestine<sup>84</sup> | RNAi: 12% embryonic lethal<sup>83</sup> |
|  | LP1064 | 99% |  |  |
| *wshc-5* | LP902 | 100% | Localizes to vesicle-like structures in the cytoplasm of *C. elegans* embryos.<sup>37</sup> | RNAi: at least 10% embryonic lethal<sup>85</sup> |
| *unc-34* | APL38 | 99% | Localizes to apical epidermal junctions and leading edge of cells during ventral enclosure in immunostained *C. elegans* embryos<sup>46</sup> | No known embryonic phenotype |
|  | LP1084 |  |  |  |
| *cap-1* | LP888 | 100% | Localizes to cell-cell contacts and to vesicle-like cytoplasmic structures in *C. elegans* embryos<sup>73</sup> | RNAi: 47% embryonic lethal<sup>86</sup> |
| *cap-2* | LP1059 | 99% | Colocalizes with F-actin at cell cortex and lamellipodia in interphase HeLa cells; colocalizes with F-actin at cortex and cleavage furrow in dividing cells<sup>87</sup> | RNAi: 46.42% embryonic lethal<sup>88</sup> |
|  | LP1060 | 99% |  |  |
| *fli-1* | LP897 | 100% | Transgenic FLI-1::GFP colocalizes with F-actin in *C. elegans* body-wall muscle<sup>89</sup> | *fli-1(bp130)* mutant is 32.3% embryonic lethal; Mutants have defects in 1st cell division<sup>89</sup> |
| *viln-1* | PHX9966 | 100% | Colocalizes with F-actin in podosome cores and at subcortical actin filaments in immunostained macrophages<sup>90</sup> | No known embryonic phenotype<sup>91</sup> |
| *add-2* | LP1032 | 100% | Localizes to the leading edge of lamellipodia in migrating MDCK cells<sup>92</sup> | No known embryonic phenotype |
| *unc-94* | LP895 | 100% | Localizes to the pointed ends of thin filaments and to muscle cell boundaries by immunofluorescence in adult *C. elegans*<sup>93</sup> | RNAi enhanced lethality from 80% to 100% in *hmp-1(fe4)* background, but no lethality on its own<sup>94</sup> |
| *gmf-1* | LP1065 | 100% | Localizes to the nucleus and co-localizes with F-actin and cofilin at membrane ruffles at the leading edge of BAECs and HMVECs<sup>95</sup> | No known phenotype |
|  | LP1076 | 100% |  |  |
| *pod-1* | LP960 | 100% | Cortical and at cell contacts in immunostained early *C. elegans* embryos, asymmetric towards anterior<sup>96</sup><br>Colocalizes and polarizes with Arp2/3 in *C. elegans* zygotes<sup>97</sup> | RNAi: 100% embryonic lethal<sup>96</sup> |
| *unc-60a* | LP1067 | 99% | Localizes to cell contacts and diffuses in cytoplasm in early *C. elegans* embryos by immunofluorescence<sup>31</sup> | RNAi: over 50% embryonic lethality<sup>31</sup> |
| *aipl-1* | LP919 | 100% | Endogenously tagged AIPL-1::GFP is detectable at very low levels in wild-type adult body-wall muscle<sup>98</sup> | *aipl-1* and *unc-78* double null is embryonic lethal<sup>99</sup> |
| *unc-78* | PHX10008 | 99% | UNC-78 is diffuse in body-wall muscle at the 1.5-fold stage and colocalizes with actin in a striated pattern in adult *C. elegans* muscle by immunofluorescence<sup>100</sup> | *aipl-1* and *unc-78* double null is embryonic lethal<sup>99</sup> |
| *cas-1* | LP1057 | 100% | Partially colocalizes with actin and regulates actin turnover in the presence of UNC-60B in immunostained *C. elegans* muscle cells<sup>101</sup> | Homozygous *cas-1(ok1523)* was mostly larval lethal<sup>101</sup> |
|  | LP1058 | 99% |  |  |
|  | LP1077 | 100% |  |  |
|  | LP1078 | 99% |  |  |
| *twf-2* | LP923 | 99% | TWF-2 localizes to the spermathecal cortex in a spectrin-dependent manner<sup>86</sup> | No known embryonic phenotype |
| *pfn-1* | PHX10152 | Homozygous lethal | Cytoplasmic in immunostained early embryos<sup>20</sup> | RNAi: 100% embryonic lethality<sup>20</sup> |
| *tth-1* | LP1079 | 100% | Enriches at cell contacts at 2-4 cell stage and colocalizes with actin, becomes weaker until localizing to nerve ring at comma stage in immunostained *C. elegans* embryos<sup>102</sup> | Homozygous *tth-1* (gk43) died as late larvae (L3 or L4)<sup>102</sup> |
|  | LP1080 | 100% |  |  |
| *phac-1* | LP1037 | 100% | PHAC-1::wrmScarlet shows weak, widespread expression in pharyngeal and head muscles, excretory cells, and neurons projecting to the nerve ring in adult *C. elegans*<sup>103</sup> | RNAi: zygotes have enlarged PAR-2 domain that is corrected prior to cell division<sup>104</sup> |
| *fln-1* | PHX5774 | 100% | Transgenic expression colocalizes with F-actin in somatic gonad<sup>105</sup> | *fln-1(tm545)* hermaphrodites produce abnormally shaped embryos, some of which fail to hatch due to ovulation defects that damage the eggs<sup>105</sup> |
| *plst-1* | RZB213 | 99% | Localizes to cell contacts and some cytoplasmic puncta, polarized anteriorly and colocalizes with LifeAct in early *C. elegans* embryos with endogenous tag<sup>75</sup> | 51% embryonic lethal in *plst-1(tm4255)*<sup>75</sup> |
| *dbn-1* | LP954 | 100% | Localizes to basolateral membranes of *C. elegans* intestine<sup>84</sup> | 0% embryonic lethality within 24hrs of RNAi<sup>88</sup> |
| *cpn-1* | LP957 | 100% | Transgelin-2 localized to leading edge of lamellipodia; colocalized with F-actin in podosomes in BMDCs<sup>106</sup> | RNAi: early cytokinesis defects and mitotic defects<sup>107</sup> |
| *frg-1* | PHX10082 | 100% | Mostly nuclear with diffuse cytoplasmic signal in immunostained embryos<sup>108</sup> | Overexpression caused defects in ventral body-wall muscle integrity<sup>108</sup> |
| *dyn-1a* | LP1072 | 97% | At midbody and furrow membrane in immunostained dividing zygotes<sup>109</sup> | Temperature sensitive *dyn-1(ky51)* have reduced brood size due to decreased egg production and embryonic and early larval lethality<sup>109</sup> |
|  | LP1073 | 96% |  |  |
| *dyn-1b* | LP1074 | 99% | DYN-1B is polarized anteriorly in zygote, localizes to cell contacts<sup>110</sup> |  |
|  | LP1075 | 100% |  |  |
| *nmy-1* | ML2540 | 99% | In epidermis during elongation in immunostained embryos<sup>111</sup> | RNAi: 20-40% embryonic lethality, reduced brood size, dumpy, uncoordinated<sup>112</sup> |
| *hum-2* | LP1041 | 100% | Localizes to apical surface of *Drosophila* follicle cell epithelium<sup>113</sup> | No known phenotypes |
| *hum-8* | LP959 | 100% | wrmScarlet::HUM-8 localizes to the cytoplasm of intestinal epithelial cells, where it displays a punctate, vesicle-like pattern, from the L1 stage through adulthood in *C. elegans*<sup>114</sup> | RNAi: sterile with germline cytokinesis defects<sup>107</sup> |
| *hmp-1* | LP191 | Published | Localizes to cell-cell contacts, with enrichment at the apical junctions of apically constricting cells in *C. elegans* embryos<sup>38</sup> | RNAi: 0% gastrulation defects, 100% embryonic lethal<sup>50</sup> |
| *afd-1* | LP650 | 100% | Localizes to cell-cell contacts and colocalizes with HMR-1/E-cadherin in *C. elegans* embryos<sup>24</sup> | *afd-1*($\Delta$), generated by introducing a premature stop codon: 23% embryonic lethality<sup>115</sup> |
| *erm-1* | BOX213 | 89% | At cell cortex of immunostained embryos, becomes apically enriched in some tissues beginning at lima bean stage<sup>116</sup> | *erm-1* RNAi causes approximately 90% early-L1 lethality, accompanied by intestinal and excretory-canal cysts<sup>117</sup> |
| *frm-1* | LP1034 | 100% | Localizes to plasma membrane in immunostained *C. elegans* embryos<sup>118</sup> | Delayed hatching - about 36% of *frm-1(tm4168)* eggs were unhatched 12 hours post-lay, reduced compared to WT<sup>118</sup> |
| *spc-1* | GOU2936 | 98% | Localizes to plasma membrane of most embryonic cells from the one-cell stage in immunostained *C. elegans* embryos<sup>119</sup> | Loss of function disrupts epidermal actin organization and embryonic elongation, causing approximately twofold larval arrest and lethality shortly after hatching<sup>119</sup> |
| *unc-70* | GOU3103 | 99% | Localizes to cell membranes in immunostained *C. elegans* embryos, but is excluded from the apical membranes of the intestine and epidermis<sup>119</sup> | 6.67% embryonic lethality within 24hrs of RNAi, 17.5% within 24-48hrs of RNAi<sup>88</sup> |
| *sma-1* | PHX4954 | 100% | Cortical dots, concentrated at cell equator of dividing 1 cell embryos, decorates actin bundles but doesn't overlap (split GFP)<sup>120</sup>In the vulva: SMA-1 is apical and UNC-70 is basolateral, SMA-1 required for SPC-1 apical localization<sup>76</sup> | *sma-1(ru18)*: 5.3% embryonic lethality<sup>121</sup> |
| *ani-1* | MDX29 | 99% | At cortex and cytokinetic ring in early embryos<sup>122</sup> | 100% embryonic lethality by RNAi<sup>122</sup> |
| *ibar-1a* | LP961 | 99% |  | No known phenotypes |
| *ibar-1b* | LP962 | 100% | Localized to apical membrane initiation site and cytokinetic bridge in C6 cells<sup>123</sup> |  |
| *pat-6* | LP958 | 100% | Body-wall muscle attachments<sup>124</sup> | RNAi: 100% paralyzed arrest at 2-fold stage (pat)<sup>124</sup> |
| *dys-1* | JIP2247 | 100% | Broad sarcolemmal signal on the outer (epidermis-facing) surface of body-wall muscle, with enrichment at the anterior tip of each muscle cell<sup>78</sup> | No known embryonic defects; exaggerated head bending<sup>78</sup> |
| *alp-1* | PHX9999 | 100% | GFP fusion protein first detected at hypodermal cell-cell junctions and in muscles during comma stage<sup>125</sup> | Small percentage of mutants had defects in actin organization, but all mutants were viable with normal brood size and muscle function<sup>126</sup> |
| *lasp-1* | LP1035 | 100% | Localizes to dense bodies and cell-cell attachment sites in body-wall muscles<sup>127</sup> | No known embryonic phenotype |
| *ehbp-1* | LP1039 | 100% | Transgene present in all somatic adult tissues in a punctate or tubular pattern<sup>128</sup> | *ehbp-1(tm2523)* mutants display larval arrest, surviving adults were sterile<sup>128</sup> |
| *hipr-1* | PHX9957 | 99% | Localized to puncta in soma and dendrites of cultured hippocampal neurons<sup>129</sup><br>Colocalized with F-actin in initiating segments of neurites<sup>130</sup> | No known embryonic phenotype; Has reduced brood size<sup>131</sup> |

**Table S4.**
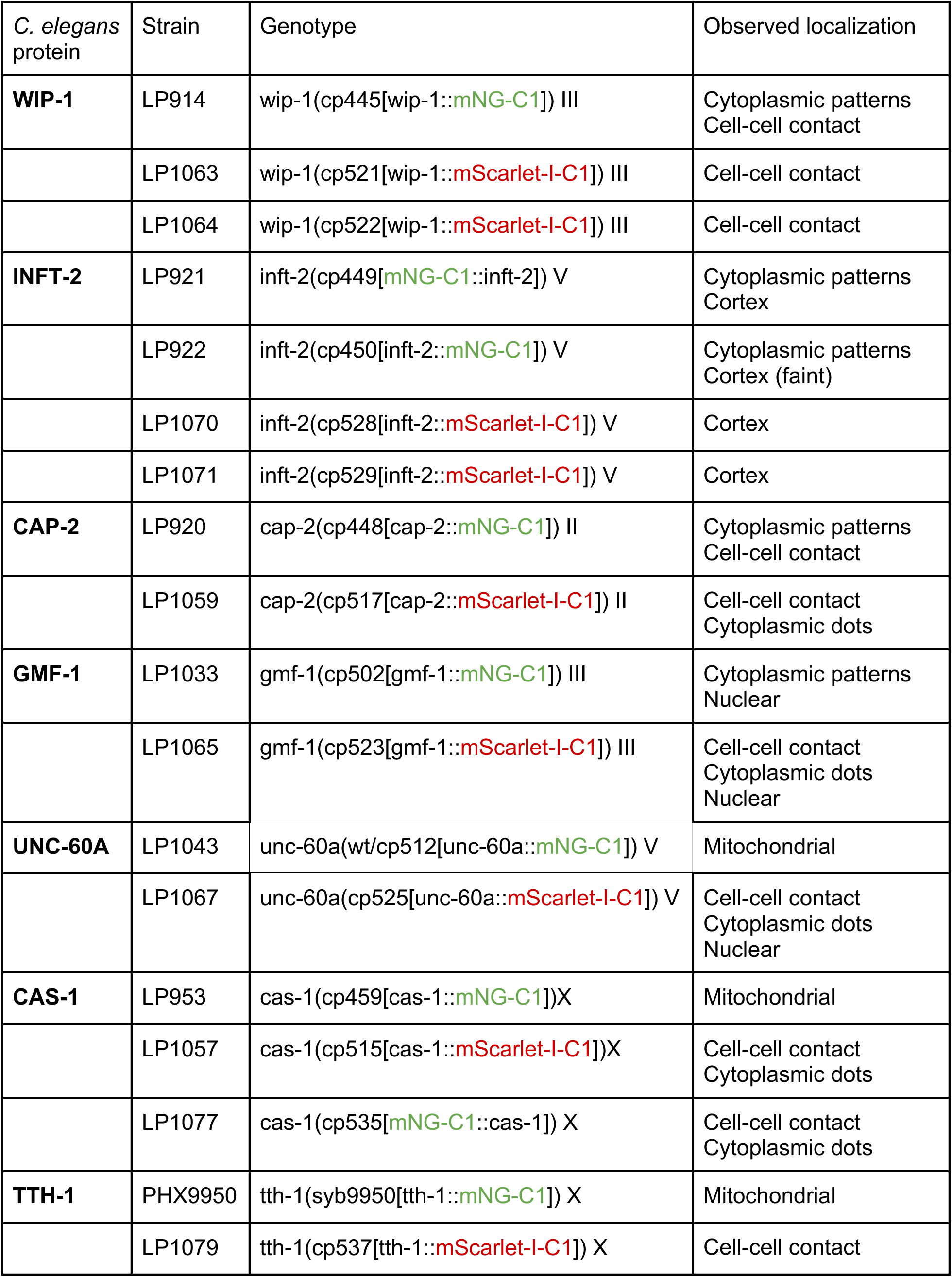
Fluorescent protein identity and tag position can affect protein localization, related to Figure 2, Figure S1 and Table 1. Summary of localization differences observed among ABPs with different fluorescent proteins and/or tag positions.

| *C. elegans* protein | Strain | Genotype | Observed localization |
| --- | --- | --- | --- |
| **WIP-1** | LP914 | wip-1(cp445[wip-1::mNG-C1]) III | Cytoplasmic patterns<br>Cell-cell contact |
|  | LP1063 | wip-1(cp521[wip-1::mScarlet-I-C1]) III | Cell-cell contact |
|  | LP1064 | wip-1(cp522[wip-1::mScarlet-I-C1]) III | Cell-cell contact |
| **INFT-2** | LP921 | inft-2(cp449[mNG-C1::inft-2]) V | Cytoplasmic patterns<br>Cortex |
|  | LP922 | inft-2(cp450[inft-2::mNG-C1]) V | Cytoplasmic patterns<br>Cortex (faint) |
|  | LP1070 | inft-2(cp528[inft-2::mScarlet-I-C1]) V | Cortex |
|  | LP1071 | inft-2(cp529[inft-2::mScarlet-I-C1]) V | Cortex |
| **CAP-2** | LP920 | cap-2(cp448[cap-2::mNG-C1]) II | Cytoplasmic patterns<br>Cell-cell contact |
|  | LP1059 | cap-2(cp517[cap-2::mScarlet-I-C1]) II | Cell-cell contact<br>Cytoplasmic dots |
| **GMF-1** | LP1033 | gmf-1(cp502[gmf-1::mNG-C1]) III | Cytoplasmic patterns<br>Nuclear |
|  | LP1065 | gmf-1(cp523[gmf-1::mScarlet-I-C1]) III | Cell-cell contact<br>Cytoplasmic dots<br>Nuclear |
| **UNC-60A** | LP1043 | unc-60a(wt/cp512[unc-60a::mNG-C1]) V | Mitochondrial |
|  | LP1067 | unc-60a(cp525[unc-60a::mScarlet-I-C1]) V | Cell-cell contact<br>Cytoplasmic dots<br>Nuclear |
| **CAS-1** | LP953 | cas-1(cp459[cas-1::mNG-C1])X | Mitochondrial |
|  | LP1057 | cas-1(cp515[cas-1::mScarlet-I-C1])X | Cell-cell contact<br>Cytoplasmic dots |
|  | LP1077 | cas-1(cp535[mNG-C1::cas-1]) X | Cell-cell contact<br>Cytoplasmic dots |
| **TTH-1** | PHX9950 | tth-1(syb9950[tth-1::mNG-C1]) X | Mitochondrial |
|  | LP1079 | tth-1(cp537[tth-1::mScarlet-I-C1]) X | Cell-cell contact |

